# CD8-mediated organization of the TCR–pMHC interface shapes its force response and dissociation pathways

**DOI:** 10.64898/2026.06.16.732622

**Authors:** Jing Li, Zhenhai Li

## Abstract

The mechanical response of the interaction between the T cell receptor (TCR) and the peptide major histocompatibility complex (pMHC) is fundamental to antigen recognition, but the atomic-scale mechanisms by which the CD8 coreceptor modulates the complex’s conformational states and force-bearing behavior remain poorly understood. We employed all-atom molecular dynamics and steered molecular dynamics simulations of membrane-embedded TCR–pMHC and TCR–pMHC–CD8 complexes to characterize their dynamics and force-induced dissociation. Microsecond-long molecular dynamics (MD) simulations show that binding of CD8 to the MHC α_3_ domain applies restraints to the latter one, which leads the MHC α_1_ helix to stably bind against the TCR complementarity-determining regions (CDRs) and suppresses the fluctuations of the antigenic peptide. Furthermore, under mechanical loading, the TCR–pMHC–CD8 system exhibits a distinct dissociation pathway compared to that of TCR–pMHC complex, which may strengthen the mechanical stability of the binding of TCR–pMHC. Collectively, these findings unravel the molecular mechanisms of CD8-mediated synergistic stabilization and mechanical regulation of TCR–pMHC, providing new mechanistic insights into coreceptor-dependent T cell antigen recognition.

**SIGNIFICANCE:** The TCR–pMHC handshake is the definitive spark that ignites the immune response. It has been shown that the force applied to the TCR–pMHC complex is critical for triggering the downstream signaling. In addition, TCR recognition of pMHC is regulated by force and strongly influenced by coreceptors, such as CD8. Therefore, understanding how coreceptors mediate TCR–pMHC interactions upon the application of force is crucial for revealing the biophysical basis of immune signaling. This study unravels how coreceptors shape the dynamics and mechanical response of the TCR–pMHC complex at the atomic level.

## 1. Introduction

The T cell receptor (TCR) precisely recognizes antigenic peptides presented by the peptide major histocompatibility complex (pMHC), which is the central event in initiating adaptive immune responses (1–3). This recognition process determines the specificity and sensitivity of the immune reaction, triggers T cell activation, and subsequently induces downstream immune cascades (4,5). Notably, TCR recognition of pMHC depends on the cooperative activity of various co-receptors, including CD4 and CD8, which play critical roles in facilitating the pMHC recognition (6–8).

In the immune system, CD8 molecules exist in two forms: the αα homodimer is predominantly expressed on natural killer (NK) cells, and the αβ heterodimer presents on about 90% of cytotoxic T lymphocytes (CTLs) (9,10). CD8αβ is essential for T cell activation through stabilization of TCR–pMHC interactions on the cell surface (11,12). As a key co-receptor, CD8 has a dual function: its extracellular domain binds to the α_3_ domain of MHC, stabilizing the TCR–pMHC interaction (13), while its cytoplasmic tail recruits the lymphocyte-specific kinases (LCKs) to initiate intracellular signaling (14,15). Existing evidence indicates that CD8 can influence TCR binding by prolonging bond lifetime (16,17).

In recent years, increasing evidence has shown that TCR recognition is strongly influenced by mechanical forces generated at the immunological synapse (18–23). Mechanical loading can regulate the lifetime and stability of the TCR–pMHC bond, induce conformational rearrangements at the binding interface, and thereby affect antigen discrimination. Experimental studies using biomembrane force probe (BFP) and optical tweezers have revealed that the interaction between TCR and agonist pMHC exhibits catch bond behavior, in which moderate forces prolong bond lifetime and enhance signaling (22,23). Under mechanical loading, the cooperative dissociation of the TCR–CD8–pMHC complex can further extend bond lifetime and enhance antigen recognition. This cooperation does not operate via a simple linear addition of bond lifetime, rather, it is governed by an allosteric conformational regulation (24–27). However, the single-molecule force spectroscopy (SMFS) struggles to reveal atomic-level details during the force-induced conformational change. To address this limitation, molecular dynamics (MD) simulations have been widely used in studies of TCR–pMHC as an assistant to bridge the gap between observations of SMFS and molecular mechanisms. The MD simulations enable the characterization of conformational change and interfacial reorganization during binding and dissociation processes. MD studies have demonstrated that TCR can adapt to different antigenic peptides through conformational selection of its CDR loops and illustrated the mechanism of cross-reactivity of TCR at atomic-level (28). Upon binding to TCR, the conformational fluctuations of both the peptide and MHC are reduced, refining the understanding of TCR–pMHC binding (29). Multiscale simulations unravel pronounced long-range allosteric coupling in this system. Such coupling is mediated by coordinated motions between structural domains, which regulate receptor dynamics and influence antigen recognition (30–33). It has been proven the TCR–pMHC interaction is strongly regulated by tensile force (34,35). Agonist pMHC interacts with TCR in a catch bond manner, while non-agonist pMHC interacts with TCR in a slip bond manner. To investigate the force-induced dissociation between TCR–pMHC, steered molecular dynamics (SMD) simulations (22,33) were carried out on several TCR–pMHC complexes with either agonist or antagonist peptides. The SMD simulations reveal distinct dissociation pathways of agonist and antagonist pMHC from TCR, which provide a molecular mechanism of the catch or slip bonds between TCR–pMHC.

Plenty of MD simulations have been carried out to investigate the dynamics and force-regulated dissociation of TCR–pMHC. However, the atomic-level mechanisms underlying CD8-mediated modulation of TCR–pMHC interactions remain unclear. To investigate the regulatory role of the co-receptor CD8 on TCR–pMHC interactions, we constructed membrane-embedded models of TCR–pMHC–CD8αβ complex. In addition, a membrane embedded TCR–pMHC was build for comparison. The bi-molecular and tri-molecular complexes will be refered by CD8– and CD8+ system in this work. Microsecond-scale MD simulations and a serial of independent SMD simulations were carried out to systematically characterize the dynamic conformational features of the complexes under force-free conditions and their dissociation pathways under pulling force. The μs-long simulations show that the binding between TCR and pMHC is highly dynamic. The binding interface shift along with Clear rotations of pMHC around TCR was observed in CD8– system. The co-receptor CD8 binding to pMHC α_3_ greatly impedes the rotation movement of pMHC around TCR, which in turn stabilizes the TCR–pMHC interface. Moreover, CD8 also influence the force-regulated dissociation of pMHC from TCR. On one hand, CD8 binding introduce additional hydrogen bonds (H-bonds), salt-bridge, and hydrophobic interactions, on another hand, CD8 binding reshapes the interaction network between TCR and pMHC, which eventually modulates the dissociation pathway under mechanical pulling. This study reveals how CD8 modulates the structural configuration and mechanical response of the TCR–pMHC complex at the atomic level. It provides a mechanistic basis for understanding co-receptor regulation in force-dependent antigen recognition.

## 2. MATERIALS AND METHODS

### 2.1 Structural Modeling of TCR–pMHC(–CD8) Complexes

The initial model was constructed based on a cryo-EM structure of the fully assembled TCR/CD3–pMHC complex and (36) (PDB: 7PHR), the crystalized structures of human pMHC-CD8αα complex (13) (PDB: 1AKJ) and mouse pMHC–CD8αβ complex (37) (PDB: 3DMM). The full-length TCR–pMHC complex model was obtained simply by removing CD3 from the cyro-EM structure of 7PHR. To specifically investigate CD8 influence on the TCR–pMHC interactions, a human TCR–pMHC–CD8 complex was built. Since the complete structure of the human CD8β remains unresolved, a human CD8β structure was predicted by AlphaFold (38). The human CD8α from 1AKJ and the predicted CD8β were structurally aligned to the mouse CD8αβ (PDB: 3DMM) to build the human CD8αβ with PyMOL (Schrodinger; New York, NY). The transmembrane (TM) segments of CD8αβ were generated by AlphaFold. Considering the existence of inter-chain disulfide bonds between α chain C143/C160 and β chain C134/C147 (39), the corresponding cysteins in the flexible linker were manually positioned in close contact allowing the formation of disulfide bonds. All remaining missing residues were modeled using Modeller (40) based on sequences obtained from the UniProt database (41). The modeled human CD8αβ and the full-length human TCR–pMHC complex were eventually aligned to the mouse TCR–pMHC–CD8αβ complex to build the human TCR–pMHC–CD8 comlex.

### 2.2 Lipid bilayer embedding and system assembly

To simulate a realistic mammalian cell membrane environment, two heterogeneous lipid bilayer systems with identical lipid compositions but different system sizes, for TCR–pMHC and TCR–pMHC–CD8 complexes, were constructed using the CHARMM-GUI Membrane Builder (42,43). The lipid bilayer was composed of 90% phosphatidylcholine (POPC) , 3% phosphatidylserine (POPS) and 7% phosphatidylinositol (4,5)-bisphosphate (PIP2) (44,45). The PIP2 was represented as an equimolar mixture of SAPI24 and SAPI25 (3.5% each), accounting for the protonation state of PIP2 at neutral pH (46,47). The protein complexes were first oriented within the lipid bilayer using the Orientations of Proteins in Membranes (OPM) database (48) and then embedded into a mixed lipid bilayer. The membrane-embedded protein complexes were placed in simulation boxes of 14.6×14.6×22.3 nm^−3^ for TCR–pMHC and 20.3×20.3×25.4 nm^−3^ for TCR–pMHC–CD8 with periodic boundary conditions. Lipid molecules were equally distributed between the upper and lower leaflets, with a total of 660 lipids for the TCR–pMHC and 1188 lipids for the TCR–pMHC–CD8 (Fig. 1). The assembled systems were solvated using the TIP3P water model, and appropriate numbers of Na^+^ and Cl^−^ ions were added to neutralize the net charge and to maintain a physiological salt concentration of 150 mM. Subsequently, the input files for MD simulations with the package of GROMACS were generated by CHARMM-GUI.

**Fig. 1.**
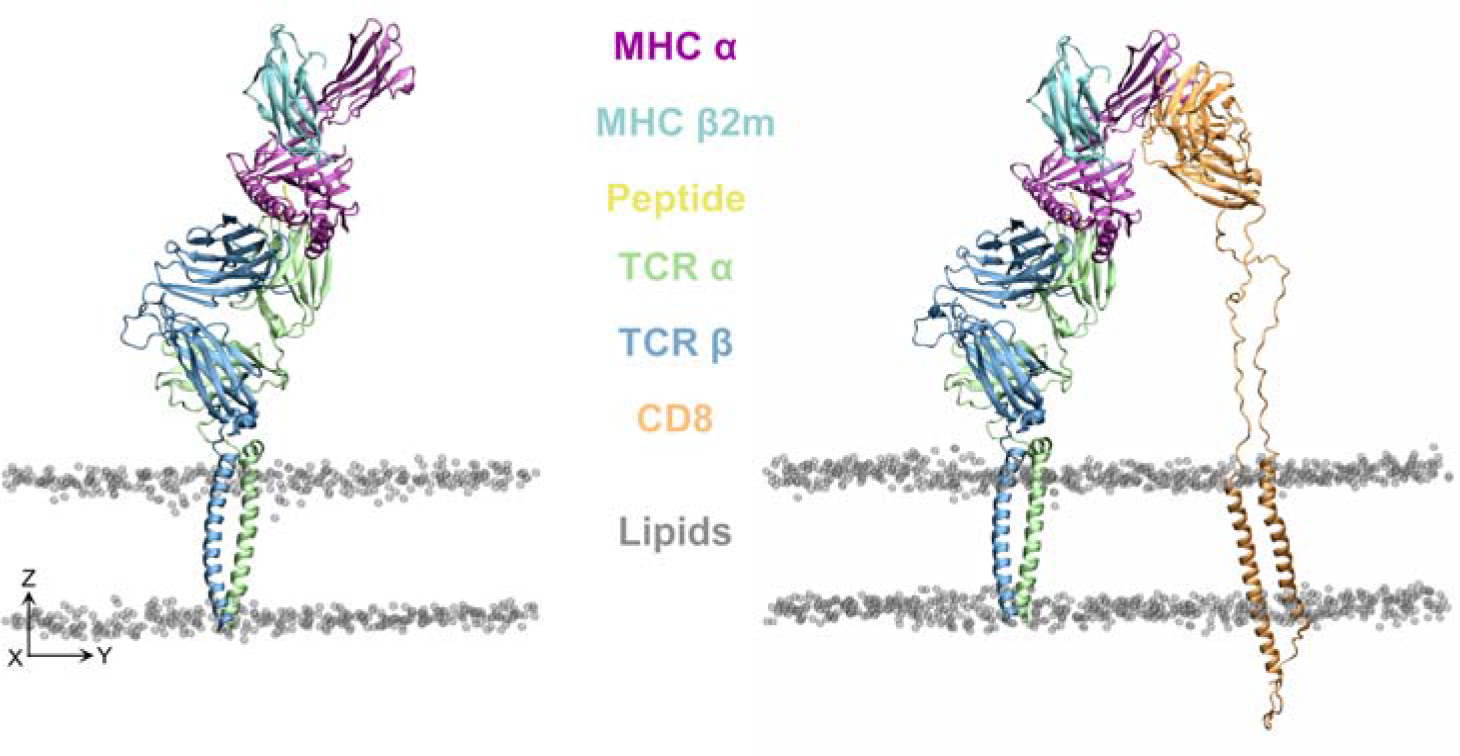
Structural models of the TCR–pMHC(–CD8) complex in a membrane environment. The molecular systems without (left) and with (right) the CD8 coreceptor were built for the following MD simulations. The TCR (α in green, β in blue), the pMHC class I complex (MHC α in purple, β2m in cyan, and peptide in yellow), and the CD8 (orange) are embedded in a realistic lipid bilayer. Gray spheres represent the phosphorus (P) atoms of the lipid headgroups, representing the membrane boundaries.

### 2.3 Equilibration and Production MD

All MD simulations were performed using GROMACS 2022 (49) with the CHARMM36 all-atom force field (50,51). Energy minimization with the steepest descent algorithm was first applied to avoid bad contacts in the system. A six-step equilibration protocol suggested by CHARMM-GUI was then applied to stabilize the systems at 310 K and 1 atm. The first two steps were conducted under the NVT ensemble for 0.25 ns with a 1 fs time step, followed by four steps under the NPT ensemble for 1.625 ns with a time step of 2 fs to adjust system pressure. Positional restraints on protein and lipid molecules were gradually released in stages until fully removed, ensuring a smooth transition of both structure and energy during equilibration.

Production MD simulations were conducted with the system temperature maintained at 310 K using the Nose-Hoover thermostat (52) (τ_T_ = 1.0 ps) and the pressure controlled at 1 atm via the Parrinello-Rahman barostat (53) (τ_p_ = 5.0 ps) with an isothermal compressibility of 4.5×10^−5^ bar^−1^. Electrostatic interactions were calculated using the particle mesh Ewald (PME) method (54), with a cutoff distance of 1.2 nm. Van der Waals interactions were treated using the same cutoff. During these production runs, a positional restraint with a force constant of 1000 kJ·mol^−1^·nm^−2^ was applied to the Cα atom at the C-terminal of the MHC heavy chain along the *z*-axis to mimic the restraints to MHC from the antigen presenting cell membrane.

### 2.4 Steered molecular dynamics

We evaluated the Root Mean Square Deviation (RMSD) of the protein in the free dynamics simulations (Fig. S2). Several structures with different RMSD were extracted from the MD trajectories (30, 50, 60, 70, and 80 ns for the TCR–pMHC system and 50, 60, 70, and 80 ns for the TCR–pMHC–CD8 system). The structures were used as the initial structures for the subsequent SMD simulations. To accommodate the dissociation distance, the simulation box for each sampled configuration was extended by 20 nm along the *z*-axis. The expanded systems were solvated and neutralized with Na^+^ and Cl^−^ ions. These systems underwent energy minimization and a six-step equilibration protocol following the same settings as described previously. An additional 10-ns MD simulation was performed for further relaxation. SMD simulations were performed using a constant velocity pulling scheme. Positional restraints with a force constant of 1000 kJ·mol^-1^·nm^-2^ were applied along the *z*-axis to the phosphorus (P) atoms of selected POPC molecules to maintain lipid bilayer stability. For the TCR–pMHC–CD8 system, a total of 64 POPC P atoms (32 per leaflet) equally distributed at the edge of the box were restrained, while for the TCR–pMHC system 48 P atoms (24 per leaflet) were restrained. The umbrella pull methods was used in SMD. A spring constant of 50 kJ·mol^-1^·nm^-2^ and a pulling speed of 0.5 nm/ns were applied to the Cα atom at the C-terminal of the MHC heavy chain. The pulling direction is perpendicular to the lipid bilayer (along the *z*-axis) to drive complex dissociation.

### 2.5 Structural Analysis

The relative rotations between TCR and pMHC were quantified using three characteristic angles, the rolling angle (α) , the pitching angle (β), and the torsion angle (γ) (Fig. 2a, b). The rolling angle and the pitching angle characterize rotational motion of the MHC α_1_-α_2_ domains relative to TCR along two perpendicular geometric directions (Fig. 2a, b), while the torsion angle represents the twisting of MHC relative to TCR along the pulling axis (Fig. 2a). To quantitively calculate these angles, two reference vectors of MHC (Line 1 and Line 2) and a reference plane of TCR (Surface 1) were defined. Line 1 is defined by the axis of the MHC α□ helix from centroids of residues 73-84 to 58-69. Line 2 is defined along the line connecting the α□ and α□ helix centroids, pointing toward α□ (Fig. S1a). The TCR reference plane was defined by three Cα atoms of variable region: Vα-W35, Vβ-W35, and Vβ-F100 (Fig. S1b). The rolling angle (α) is defined as the angle between Line 1 and the normal vector of the TCR reference plane, whereas the pitching angle (β) is defined as the angle between Line 2 and the normal vector of the TCR reference plane. The torsion angle γ is defined by the angle between the projection of Line 1 onto the TCR reference plane and the vector in the plane from Vβ-W35 to Vα-W35.

**Fig. 2.**
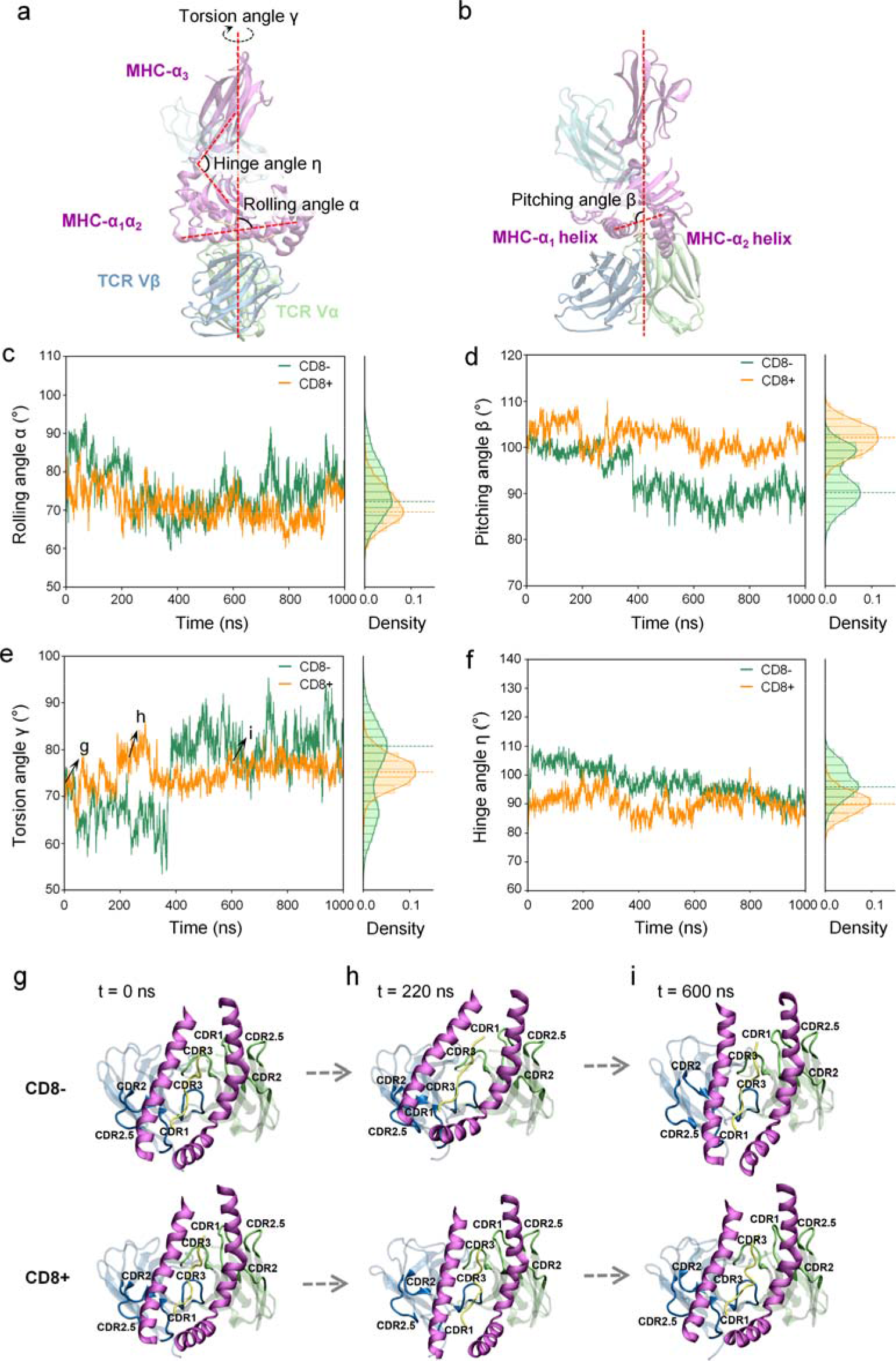
Geometric definitions of TCR–pMHC orientations and their dynamic stability in the force-free MD simulations. a, b. Geometric definitions of the rolling angle α, pitching angle β, torsion angle γ, and hinge angle η (details provided in the Methods). c-f. Time-course plots (left) and distributions (right) of angles α, β, γ and η during 1,000 ns of force-free MD simulations. Dashed lines in the density histograms indicate the peak positions of the angle distributions in the force-free MD simulations. g-i. Representative snapshots of CD8– (upper) and CD8+ (lower) systems at t = 0, 220, and 600 ns illustrate the rotational rearrangements of pMHC relative to TCR corresponding to changes in the torsion angle γ.

In the simulation, rotation of α_1_ and α_2_ domains around α_3_ domain was observed during pulling applied. To quantify the rotation, two vectors, one in the α□-α□ platform and the other in the α□ domain, were used. The vector in α□-α□ is from the centroid of residues S11, V12, F22, I23, V95, and Q96 to that of residues Y7, F8, G26, Y27, Y99, and G100, while the vector in α_3_ is from the centroid of residues R202, C203, E229, L230, A246, and V247 to that of residues S207, F208, P235, A236, F241, and Q242. The angle η between these two vectors was calculated to evaluate the conformational change of MHC (Fig. S1c). The geometric analyses mentioned above and structural visualization were carried out using the package, Visual Molecular Dynamics (VMD) (55).

## 3. Results

### 3.1 CD8 stabilizes the TCR–pMHC interface in the force-free condition

The human TCR–pMHC–CD8 complex structure is constructed based on a human TCR–pMHC complex (36), an AlphaFold-modeled CD8 (38), and a mouse pMHC–CD8 complex (37). Then the modeled TCR–pMHC–CD8 and TCR–pMHC complexes were inserted into a heterogeneous lipid bilayer to construct the membrane-embedded protein complexes (see details in Materials and Methods, Fig. 1). The membrane-embedded complexes were soaked in neutralized NaCl solution with a concentration of 150 mM to mimic the physiological condition. Then, 1 μs-long free MD simulations were performed on both systems with the edge of the lipid bilayer restrained. As in the physiological condition, the C-terminus of the MHC α chain is inserted into the antigen-presenting cell membrane, the position restraint in the z-direction was introduced to the C-terminal of the MHC α chain.

In the μs-long simulation trajectories, clear fluctuations were observed in the protein complex. To pinpoint the source of the fluctuation, the RMSD traces of the whole complex, TCR alone, and pMHC alone were calculated, respectively. The significant and moderate oscillations were respectively found in the RMSD of the whole complex (Fig. S2a) and pMHC alone (Fig. S2a), especially in the CD8– system. RMSD of TCR alone keeps relatively stable in the early stage but shows a noticeable increase around ∼400 ns (Fig. S2c). In contrast, the tri-molecule complex exhibited a relatively stable dynamic profile for the whole complex, pMHC alone, and TCR alone (Fig. S2; Supplementary Video S1). Given that the internal conformational fluctuation within TCR and pMHC alone were subtle compared to the whole complex, the RMSD oscillations of the whole complex primarily arise from the relative motion between molecules. To further validate this speculation, we calculated the interface root-mean-square deviation (iRMSD) to quantify the inter-molecular movement (Fig. S2d). The TCR variable domains, the MHC α_1_-α_2_ domains, and the antigenic peptide are all included in the interface. The iRMSD of CD8– system exhibits multiple metastable states, and the molecule complex hops back and forth within these states, whereas the CD8+ is relatively stable with a single state.

Given the fact that the pMHC molecule stably sticks to the TCR variable region throughout the free dynamics simulations, the fluctuation of iRMSD is dominantly due to the rotation of pMHC around TCR. The inter-molecular rotations were further characterized by three parameters: a rolling angle (α) and a pitching angle (β), reflecting the tilting of the MHC helices over the TCR, and a torsion angle (γ) representing its twisting around the pulling axis (Fig. 2a, b). Since the TCR–pMHC complex is less stable, the variations of the rolling, pitching, and torsion angle of TCR–pMHC complex are more pronounced than those of TCR–pMHC–CD8 complex (Fig. 2c-e). The rolling angles of both complexes gradually change without any sudden jump (Fig. 2c; Supplementary Video S2), whereas the pitching angle in the CD8– system undergoes an abrupt conformational transition at ∼400 ns (Fig. 2d; Supplementary Video S3). A careful angle distribution analysis reveals that the TCR–pMHC complex distributes broadly in rolling angle, and exhibits two peaks in pitching, indicating the existence of two metastable states. In contrast, the TCR–pMHC–CD8 complex remains more tightly constrained, with both rolling and pitching angle fluctuating within relatively narrow ranges centered around 70° and 100°, respectively (Fig. 2c, d; Supplementary Videos S2, S3).

The torsion angle γ exhibited even more pronounced differences in dynamic behavior between the two systems. In the TCR–pMHC–CD8 complex, although a transient angle increase from 75° to 80° occurs at ∼200 ns, the system rapidly relaxes back, maintaining a clear unimodal distribution centered at ∼75° (Fig. 2e, g-i; Supplementary Video S4). In contrast, the system lacking CD8 exhibited obvious conformational instability, accompanied by two torsion angle transitions. Around the initial 40 ns of simulation, γ slightly decreased to approximately 60° (Fig. 2e, g, h). At ∼400 ns, γ abruptly increased over 80° and then underwent sustained extensive fluctuations at the elevated state (Fig. 2e, i; Supplementary Video S4). As a result, in the absence of CD8, the torsion angle clearly showed a double peak in the distribution (Fig. 2e). Since the abrupt jumps of pitching and torsion both occur at the same moment, the two angles are closely correlated.

Besides the iRMSD, the RMSD of the pMHC greatly increased to approximately 3-4 Å, suggesting a moderate conformational change (Fig. S2b). By carefully examining the trajectory, a clear opening of the hinge between the α_1_-α_2_ platform and the α_3_ domain was observed. Therefore, an angle η of the hinge was defined to quantitatively characterize the geometric evolution of the MHC molecule (Fig. 2a). In free dynamic simulations, the TCR inserts into the lipid bilayer and the C-terminal of MHC α chain is restrained in the z-direction. The random diffusion of the two ends of the complex in the xy-plane may enlarge the distance between the ends. It has been reported that pulling the TCR–pMHC complex induces the opening of MHC (22). As expected, with the two ends restrained in z-direction, the hinge of MHC opens by ∼20° in the TCR–pMHC complex. In contrast, CD8 binding markedly constrained the overall geometric flexibility of the MHC, stabilizing the MHC α_3_ domain nearly perpendicular to the α_1_-α_2_ platform. As a result, the hinge angle is greatly narrowed without any significant elevation (Fig. 2f; Supplementary Video S5). These findings suggest that CD8 binding effectively anchors the MHC to TCR, imposing a robust constraint that stabilizes the binding orientation and MHC conformations.

### 3.2 CD8 impedes TCR–pMHC interface sliding and stabilizes the peptide in MD

To evaluate the contribution of CD8 to interacting residue pairs, we calculated the average distance maps between the TCR CDR loops and the MHC helices during the simulations (Fig. 3a-d). The V regions of both α and β chains make contact with MHC helices. The CDR1, CDR2, and CDR2.5 loops of the β chain form stronger interactions with the α_1_ helix, while the CDR3 loop mostly interacts with the α_2_ helix. In contrast, the CDR1, CDR2, and CDR2.5 loops of the α chain form stronger interactions with the α_2_ helix, while the CDR3 loop mostly interacts with the α_1_ helix. CD8 impedes the rotation of MHC around TCR, which could stabilize the TCR–pMHC binding interface. Direct subtraction of the distance maps of CD8–system from the other system (Fig. 3e, f) reveals binding interface shifts toward α_1_ helix in the TCR–pMHC–CD8 system, indicated by the shortened distances from all the CDR loops to the pMHC-α_1_ helix and the slightly enlarged distance to pMHC-α_2_ helix.

**Fig. 3.**
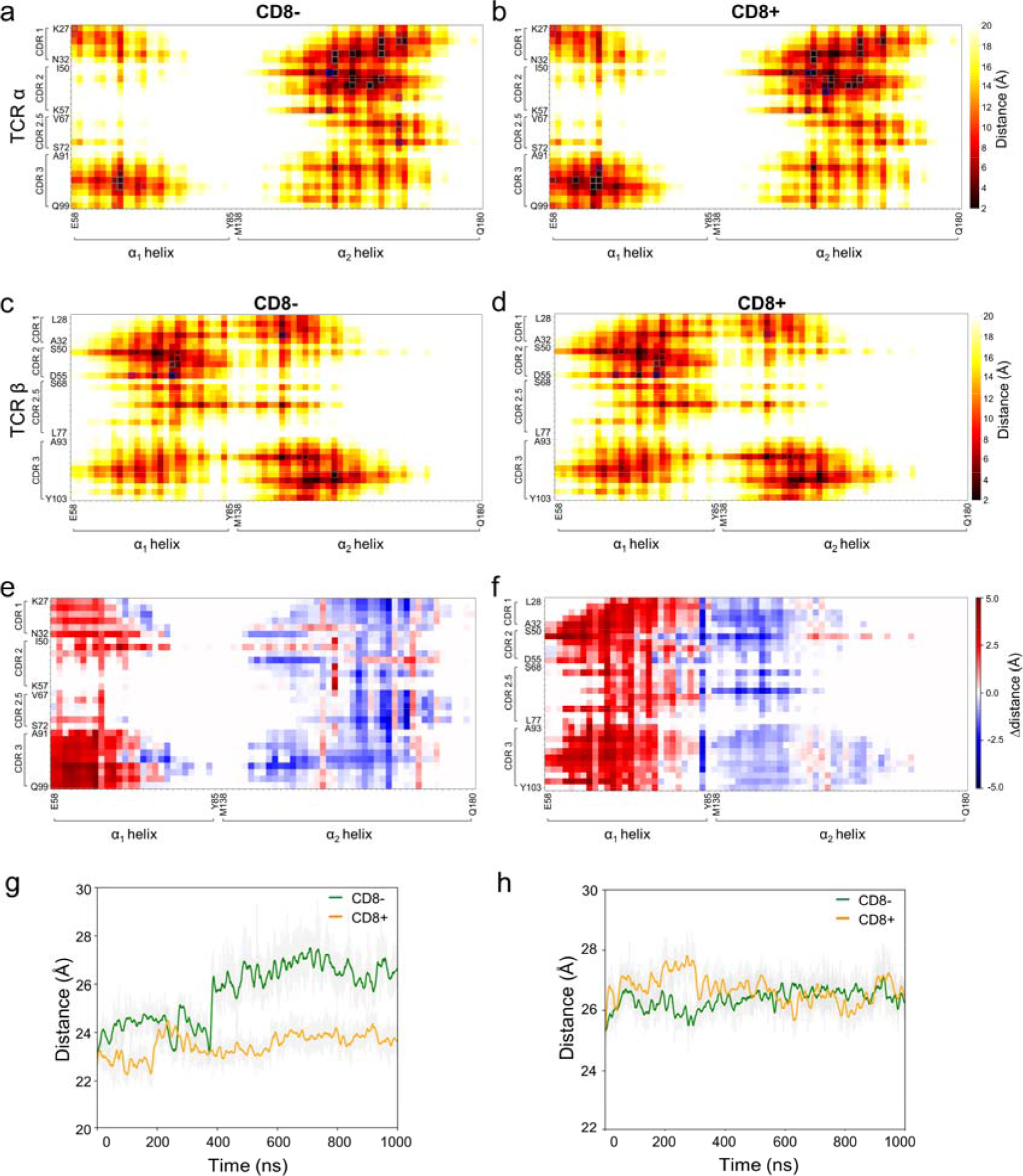
CD8 stabilization of the TCR–pMHC interfacial dynamics during force-free MD. a-d. Average contact maps between TCR α CDR loops and MHC helices in the absence (a) and presence (b) of CD8. c, d. Average contact maps between TCR β CDR loops and MHC helices in the absence (c) and presence (d) of CD8. Specific interaction types are indicated by colored outlines in the grid: gray for hydrogen bonds, red for hydrophobic interactions, and blue for salt bridges. e, f. Difference of average contact distances from the TCR α chain (e) and β chain (f) to MHC, showing changes in distances between MHC helices and TCR CDR loops; red indicates shorter distances, and blue indicates longer distances in the presence of CD8. g, h. Time evolution of the center of mass distances between the TCR V domain backbone and the MHC α_1_ (g) and α_2_(h) helix backbones.

Furthermore, we calculated the center of mass (COM) distance between the MHC α_1_/α_2_helix backbone and the TCR V-domain backbone (Fig. 3g, h). In consistent with the distance maps, the COM distance between the TCR V-domain backbone and the MHC α_1_ helix backbone jumps from ∼24 Å to ∼27 Å at ∼400 ns, the exact moment when the pitching and torsion angles suddenly hop (Fig. 3g), suggesting the interface shifting is accompanied with the rotation of pMHC relative to TCR. In contrast, the COM distance for the MHC α_2_helix shows only minor fluctuations, suggesting that its interaction with the TCR is less affected by CD8 (Fig. 3h). Moreover, we calculated the minimal distance of each residue pair at the interface of α_1_ helix to the CDR loops of both TCR chains throughout the simulation trajectory (Fig. 4 and Fig. S3). The minimal distance traces clearly reveal abrupt shifts at two specific time points, t_1_ and t_2_. At t_1_, the contact interface drifts toward the C-terminus along the α_1_ helix, while at t_2_, it reverts toward the N-terminus, which is synchronized with the hopping of the pitching and torsion angles. Notably, the contact between LEU76 and TCR β chain is constitutively maintained throughout the simulation of CD8– system. Given the corresponding decrease in torsion angle γ at t_1_ and its subsequent increase at t_2_ (Fig. 2e), the contact interface drift is attributed to the rotation of pMHC around the z-axis passing through the LEU76 anchor. In contrast, binding of CD8 stabilizes the TCR–pMHC complex and minimizes the rotations. Therefore, the drift of the interface along α_1_ helix was not observed in CD8+ system.

**Fig. 4.**
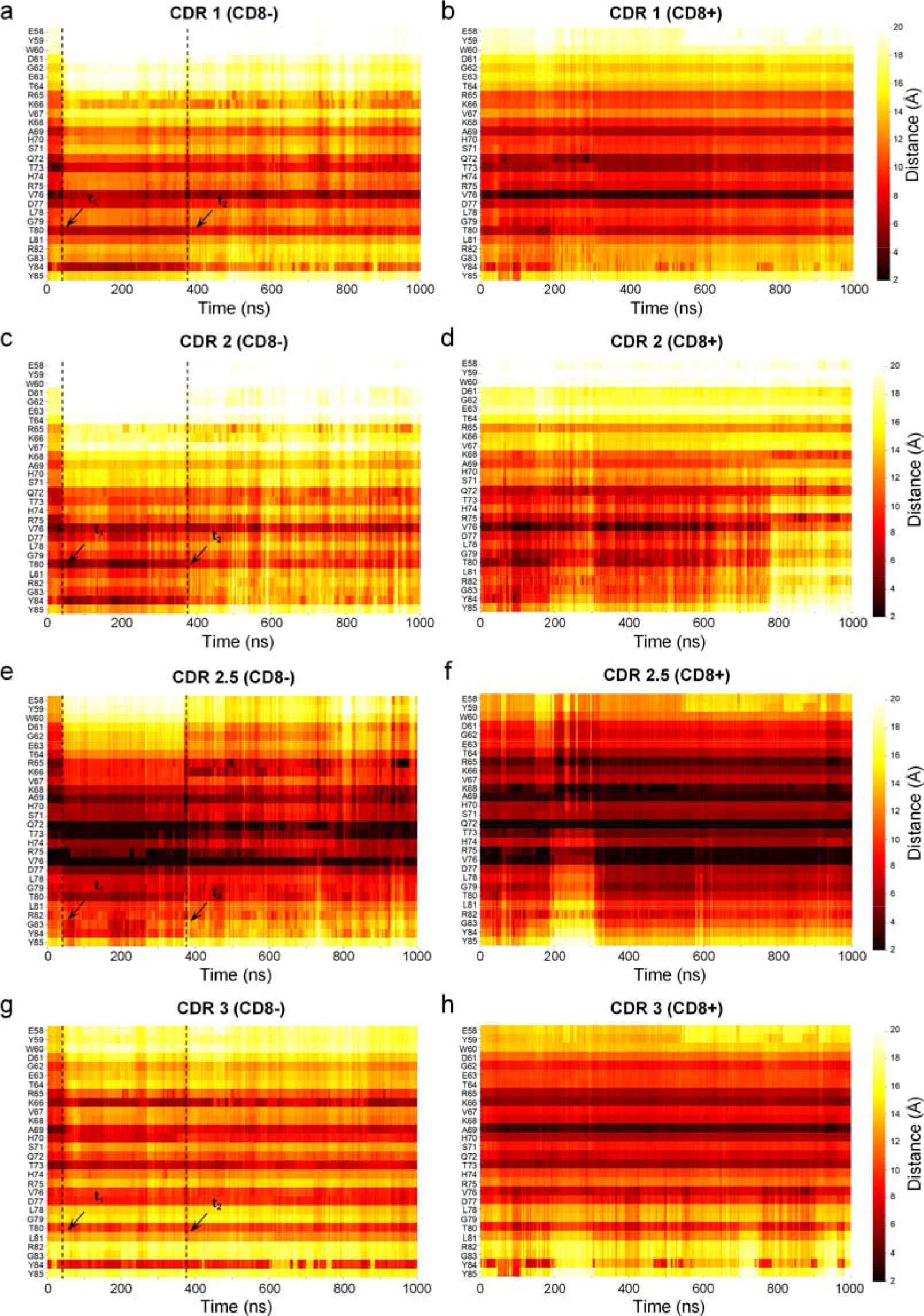
Time-resolved contacts between TCR β CDR loops and the MHC α_1_ helix during Force-free MD simulations. a-h. Time courses of minimum distances between MHC α_1_ helix residues and TCR β CDR1 (a, b), CDR2 (c, d), CDR2.5 (e, f), and CDR3 (g, h) loops in the TCR–pMHC system (a, c, e, g) and TCR–pMHC–CD8 system (b, d, f, h). Time points *t*_1_ and *t*_2_ mark interface sliding events associated with γ angle shifts.

The contact region drift can further affect the salt bridge, hydrophobic, and hydrogen bond (H-bond) formation. Therefore, we quantified the interaction occupancy for each individual TCR interfacial residue to locate the key residue for recognition with or without CD8. As expected, marked differences were observed in recognition hotspots (Fig. 5a-d and Fig. S4). At the loop level, the presence of the CD8 co-receptor differentially modulates the hydrogen-bonding networks across the TCR CDR loops. Within the TCR α chain, CD8 binding to the pMHC induces a slight increase in hydrogen bonds within the CDR1 loop, a pronounced enrichment in the CDR3 loop, and a concomitant decrease in the CDR2 loop. Conversely, the TCR β chain exhibits a distinctive response, characterized by a minor weakening of hydrogen bonds in the CDR1 loop alongside a significant structural enhancement in the CDR3 loop. The total salt bridge was slightly reduced upon binding of CD8. The basic residue R51 in α CDR2 dominates the salt bridge in the absence of CD8, while it vanishes in the presence of CD8. As an alternative, two acidic amino acids, E54 and D55, form the most salt bridges (Fig. 5b and Fig. S4a). Even though the salt bridge on R51 is suppressed, the H-bond formation of it is maintained in the presence of CD8. Noteably, Residues S29, N31, and N32 participate in transient hydrogen bonding with the pMHC under baseline conditions without CD8. Co-receptor binding alters this local network, triggering a sharp decline in the contact probabilities of S29 and N32. Remarkably, this loss is fully compensated for by a dramatic escalation in bonding frequency at N31. (Fig. 5a, d). Overall, the binding of CD8 led to a substantial increase in the total H-bonds across the entire TCR interface (Fig. 5c), suggesting that CD8-mediated stabilization not only refines specific recognition hotspots but also enhances the overall connectivity and stability of the TCR–pMHC network of hydrogen bonds.

**Fig. 5.**
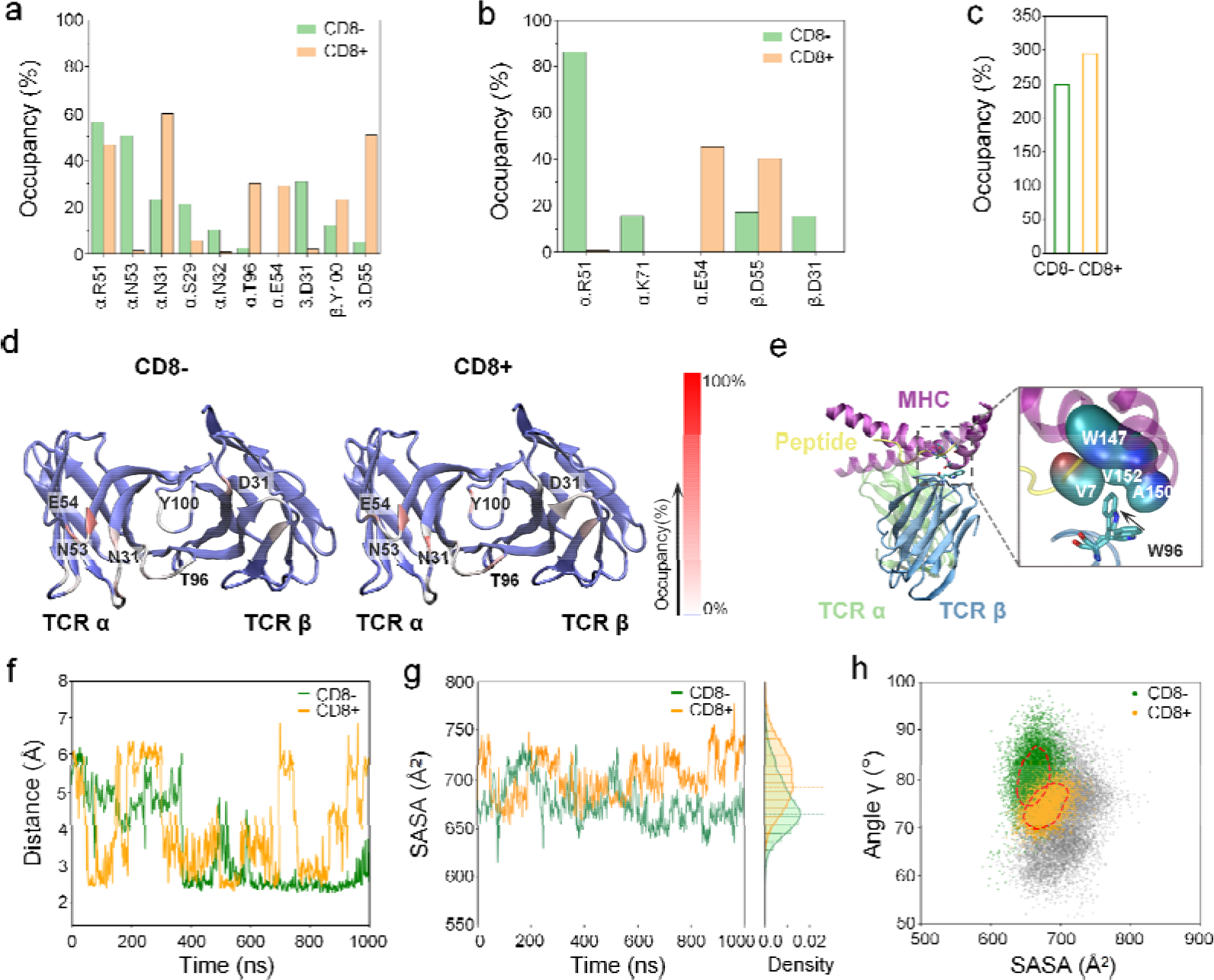
CD8-mediated TCR interactions interface. a, b. Comparison of the occupancies of H-bonds (a) and salt bridges (b) for key TCR interfacial residues between systems in the absence and presence of CD8. c. Cumulative occupancy of H-bonds formed by all participating residues on the TCR interface. d. The heatmaps of H-bond occupancy on the TCR V domains in the absence or presence of CD8. e. Schematic of the hydrophobic pocket (formed by the peptide and the MHC α_2_ helix), with the inset displaying the spatial arrangement of residues W147 (MHC), V7 (Peptide), V152, and A150 (MHC) relative to the TCR residue W96; arrow highlights the movement of residue entry into the pocket. f. Time-course of the distance between the pocket center (W147) and W96, where the dashed line at 3.2 Å marks the threshold defining the insertion into the pocket. g. Time-course (left) and density histogram (right) of SASA values for the hydrophobic pocket and the associated residues between the two systems; lower SASA values represent enhanced hydrophobic contact. Dashed lines in the density histograms indicate the peak positions of the SASA distributions in the force-free MD simulations. h. Scatter plot showing the correlation between pocket SASA and twisting angle γ; colored dots represent configurations with successful pocket entry, while gray dots represent states where the residue has not entered the pocket; dashed circle highlights the density 2% cluster.

In contrast, the residues that formed hydrophobic interactions are largely maintained, no matter whether CD8 exists or not (Fig. S4b). The amino acids, W96, A98, and P99, in β CDR3 build up a hydrophobic surface, which forms an interaction with the complementary hydrophobic pocket on MHC α_2_ helix formed by W147, A150, and V152, and peptide V7, in both cases (Fig. 5e). However, the interaction exhibits distinct dynamic evolution. Distance analysis shows that in the presence of CD8, W96 at β CDR3 steps in and out of the hydrophobic pocket during the simulation. In contrast, in the absence of CD8, the torsion of TCR around the pulling axis shortens the distance between TCR CDR loops and MHC α_2_ helix. The torsion brings the β CDR3 closer to the hydrophobic pocket and leads the W96 to insert deeply into the pocket and stably sticks around (Fig. 5f). Accordingly, a lower Solvent Accessible Surface Area (SASA) was observed in the absence of CD8 (Fig. 5g). Careful examination unravels a correlation between the SASA and the torsion angle γ, where a large torsion angle γ corresponds to a lower SASA (Fig. 5h).

Although the CD8 bound to MHC does not directly contact the peptide, it restrains the movement of the peptide. According to the Root Mean Square Fluctuation (RMSF) of the backbone (Fig. 6a) and side chains (Fig. 6d) of the peptide, the entire peptide backbone fluctuated much less significantly in the presence of CD8, while only the side chain of the N-terminal residue exhibited stronger fluctuation. Occupancy maps of the backbone (Fig. 6b, c) and sidechains (Fig. 6e, f) further show that, in the presence of CD8, the peptide adopts a more stable conformation within the MHC binding groove. In the CD8– system, the peptide residues E3, P4, and G5 exhibited higher flexibility with backbone RMSF peaks reaching ∼1.8 Å (Fig. 6a), while the sidechain fluctuations of P4 are even more pronounced, peaking at ∼2.4 Å (Fig. 6d). Therefore, we closely inspected the P4 movements in the simulated trajectories, which suggested that CD8 binding maintained the hydrophobic contacts of the peptide-P4 and MHC-A69 (Fig. 6i), while peptide-P4 flips to the other side of the groove making contact with L156 (Fig. 6h). Comparing the binding groove of CD8– and CD8+ system, a clear difference in the width of the binding groove can be observed (Fig. 6h, i). By calculating the Cα distance between A69 and L156, we quantified the size variations of the binding groove of both systems (Fig. 6g). Surprisingly, the width of the binding groove swiftly rised to 18 Å within 100 ns in the CD8– system, while it maintained 16 Å throughout the simulation with CD8+ bound to MHC. These findings indicate that binding of CD8 keeps the binding groove in a compact conformation, which may facilitate stable peptide docking through enhanced structural constraint.

**Fig. 6.**
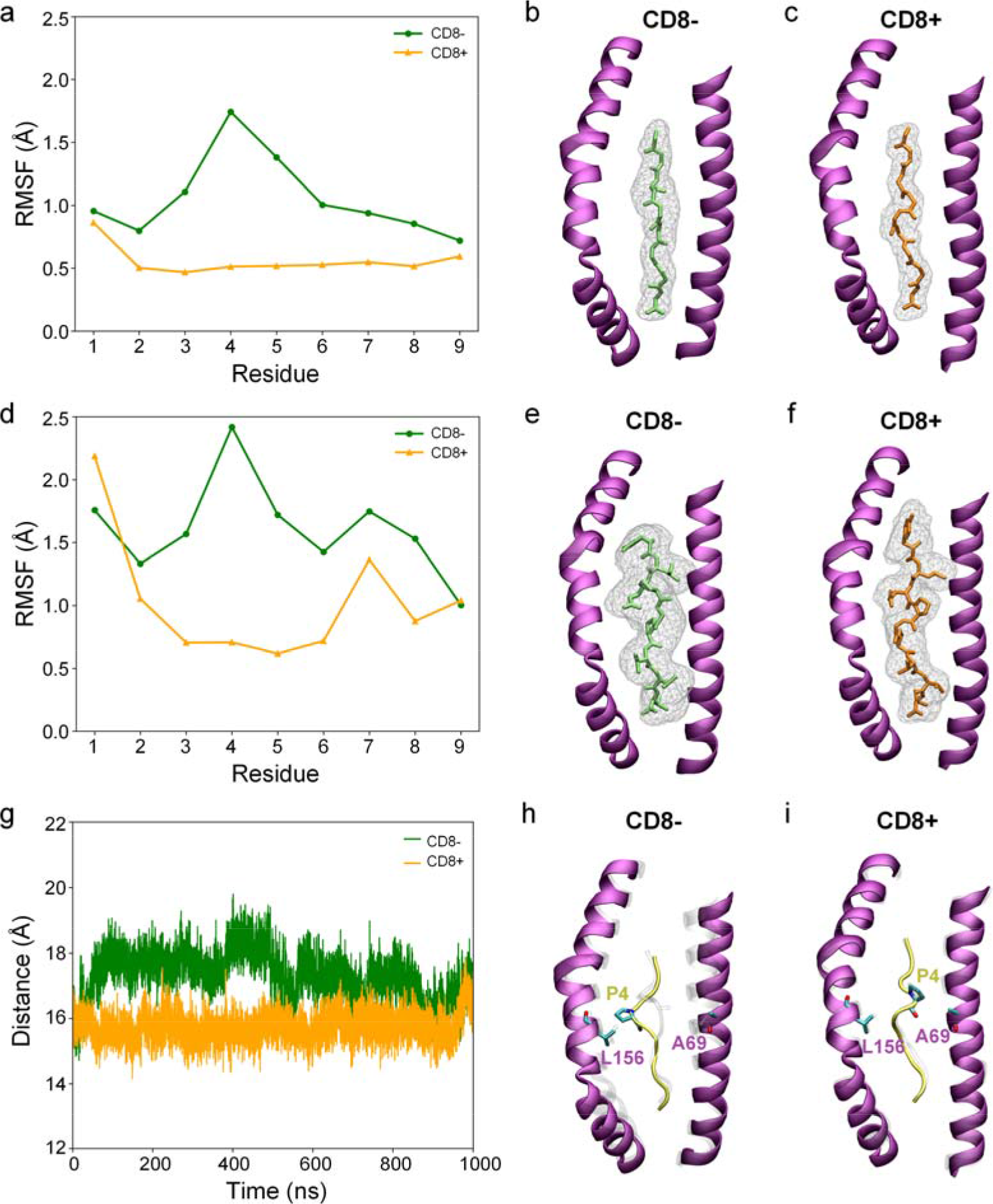
CD8 Regulation of Peptide Stability and the MHC Binding Groove. a, d. Root Mean Square Fluctuation (RMSF) of the peptide backbone (a) and sidechain atoms (d). b, c. Wireframe isosurfaces representing atomic occupancy of the peptide backbone within the MHC binding groove in the absence (b) or presence (c) CD8, calculated with VMD volmap at an isovalue of 0.03. e, f. Wireframe isosurfaces showing atomic occupancy of the peptide sidechains within the MHC binding groove in the absence (e) or presence (f) CD8, calculated and visualized as above. g. Time evolution of the Cα distance between MHC residues A69 and L156. h, i. Comparison of the peptide-P4 orientation and the groove width between A69 and L156 in the absence (h) or presence (i) of CD8. The images show overlays of simulation snapshots aligned to the crystal structure (gray).

### 3.3 Distinct dissociation pathways under pulling force

It has been demonstrated that the pulling force applied to the TCR–pMHC is critical for triggering the downstream signaling and the T cell activation (23,56). Moreover, CD8–MHC interaction can be strengthened by pulling force, which is critical for targeting cognate antigens (27). Therefore, SMD simulations were carried out to investigate the force-driven dissociation of both CD8– and CD8+ systems. To sample as many as possible dissociation pathways, multiple TCR–pMHC and TCR–pMHC–CD8 configurations with distinct RMSD obtained from the free MD simulations were used as the initial structures in the SMD simulations. In the TCR–pMHC system, all five independent trajectories of the TCR–pMHC complex underwent complete dissociation (Fig. 7a, c; Supplementary Video S6). At the beginning of the pulling, a C-terminal β-strand in the MHC α_3_ domain unfolded in 4 out of 5 simulations. Meanwhile, the TCR transmembrane domain tends to be extracted from the cell membrane. With the force increasing, the complex exhibited a continuous increase in the rolling angle α, rapidly from approximately 80° to over 100° before dissociation (Fig. 7a), indicating the rolling of TCR on the MHC interface occurs. In contrast, the torsion angle γ just fluctuated around without a clear trend. The rolling angle distribution showed that all the mean rolling angles increased by ∼3-∼10 ° and three out of five simulations exhibited sharp peaks, while the other two showed relatively broad distributions. In contrast, the torsion angle showed no systematic change across all the independent simulation trajectories. (Fig. 7c).

**Fig. 7.**
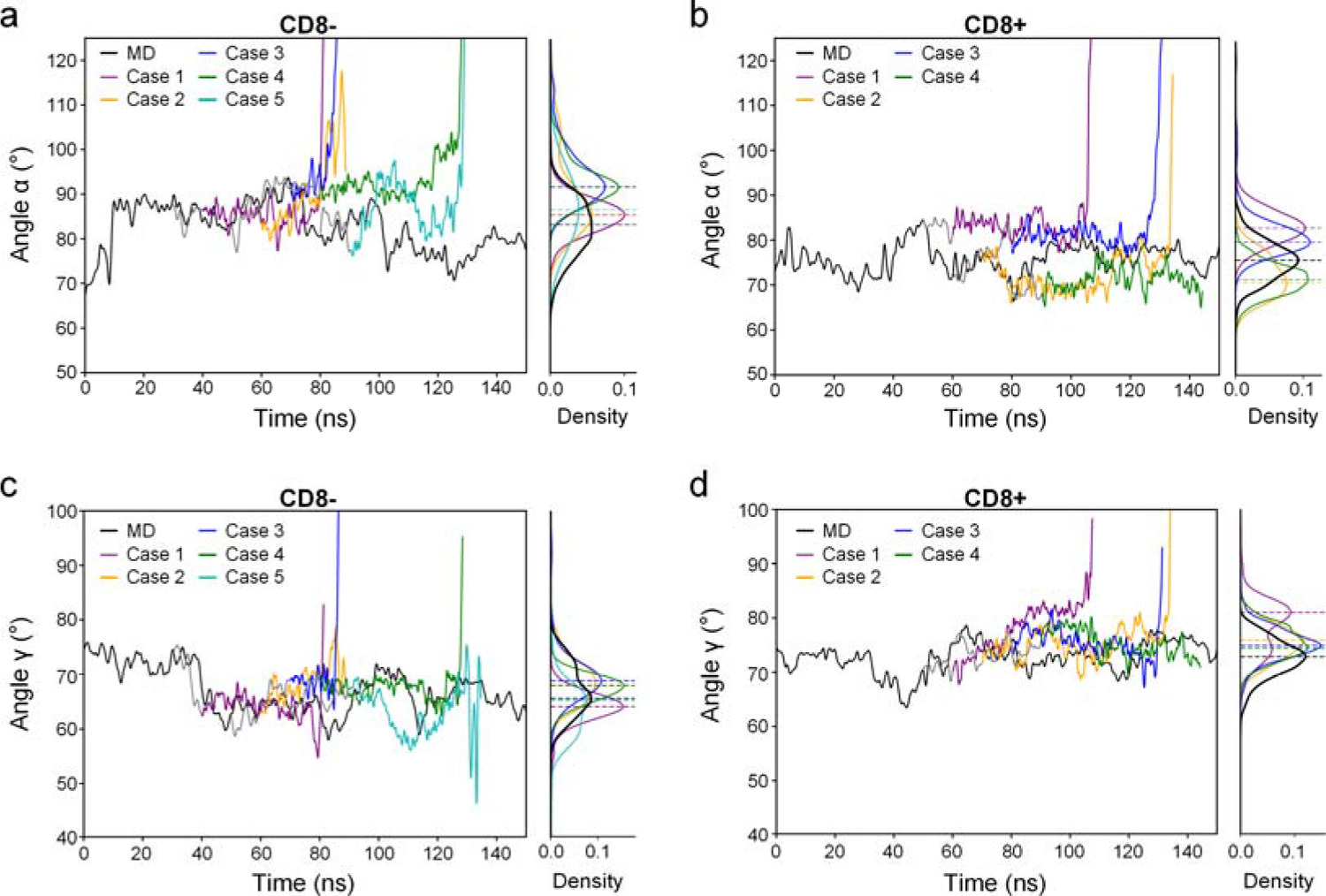
Distinct force-induced dissociation pathways in the presence and absence of CD8. Comparison of angular dynamics between MD and SMD trajectories. The rolling angle α (a, b) and the torsion angle γ (c, d) are shown for the CD8– system (a, c; five cases) and the CD8+ system (b, d; four cases). Black curves represent the force-free MD trajectories, colored curves represent multiple independent SMD trajectories from the start of pulling to complex dissociation, and gray curves represent additional 10□ns force-free MD trajectories performed after box enlargement. The density histograms (right side of each panel) of angles α, γ are calculated from all data of the free MD and SMD trajectories, and the same color code was used to distinguish different conditions. Dashed lines in the density histograms indicate the peak positions of the angle distributions for the respective runs.

In the CD8+ system, the extraction of the transmembrane domain of TCR and unfolding of the C-terminal MHC α_3_ domain were observed upon pulling (Supplementary Video S7). It’s worth noting that the α_3_ domain underwent more extensive unfolding. Partial or full unfolding of the MHC α_3_ domain was observed upon pulling. In 1 out of 4 simulation runs, with the α_3_ domain fully unfolding, the C-terminus reached the edge of the simulation box before the pMHC dissociated from TCR (Supplementary Video S7). Obviously, binding of CD8 to MHC markedly altered the force-regulated manner of TCR–pMHC (Fig. 7b, d). Under the application of external force, the rolling angle within the TCR–pMHC–CD8 complex did not consistently increase or decrease according to four independent simulations. In addition, the fluctuations of rolling angle were substantially impaired compared to those observed in free MD simulations (Fig. 7b), which suggests that mechanical tension, in coordination with CD8 binding, suppresses the rolling motion of the TCR relative to the pMHC. In contrast, the torsion angle of all four SMD simulations increased by ∼2-∼10° compared with force-free MD (Fig. 7d). Considering the mean torsion angle γ of TCR–pMHC in the CD8– system during the free MD simulation is roughly 8° higher than that in CD8+ system, the applied pulling force effectively steers the CD8-bound TCR–pMHC toward a configuration with comparable γ to that of TCR–pMHC complex alone. These results suggest that the tri-molecule complex primarily relieved external force through adjustments in the torsion angle, while the bi-molecular complex primarily relaxed through rolling. Thus, binding of CD8 to MHC shifts the dominant dissociation pathway under force from a rolling-dominated to a torsion-dominated regime.

Further structural evolution analysis revealed that this pathway shift is closely related to the dynamic reorganization of CD8. In all SMD trajectories, CD8 did not immediately detach under force but instead underwent consistent dynamic sliding along the MHC α_1_-α_2_ domains (Fig. 8a, b). In 1 out of 4 cases, the sliding eventually guided CD8 travel close to the TCR–MHC interface (Figure 8c; Supplementary Video S7). Since in the presence of CD8, the rolling angle was in a lower range, which opens the interface between TCR and pMHC (orange trajectories in Fig. 7b), enough room was provided for CD8 to wedge in. The dynamic sliding of CD8 along the MHC surface establishes a distinct load-bearing architecture that redistributes tensile stress along the TCR–pMHC axis, thereby structurally stabilizing the entire complex under load.

**Fig. 8.**
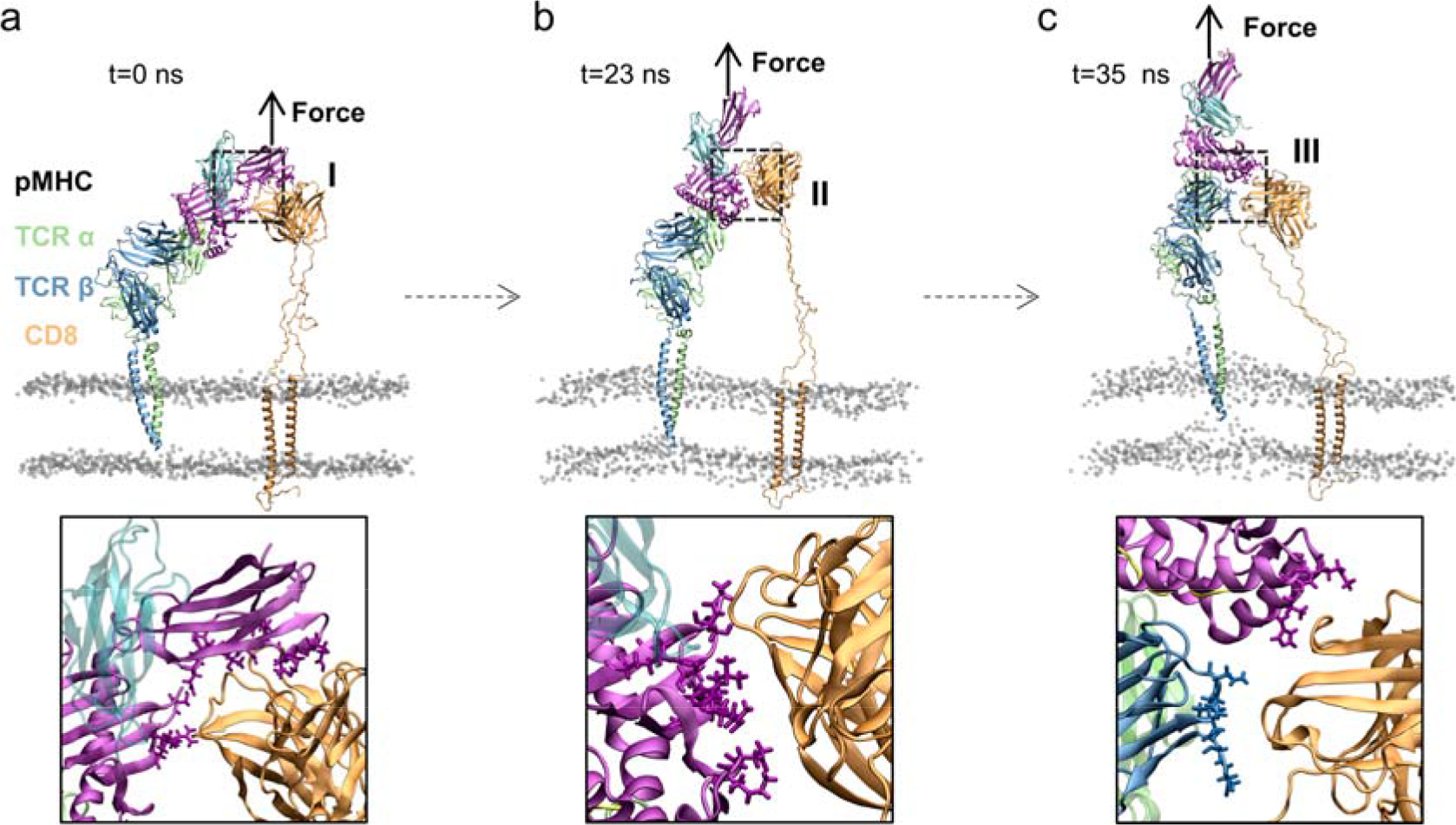
Reorganization of the CD8 binding interface under pulling force. a-c.The structure of the full complex (upper) and zoomed-in interface (lower) of the initial binding state (a), the state with CD8 sliding to the MHC α_1_-α_2_ domains (b), and with CD8 wedged into the TCR–pMHC binding interface (c). The interfacial residues within 3.2 Å of CD8 are shown.

### 3.4 Rolling driven by external force induces new interactions formed between TCR–pMHC

To investigate how external force modulates TCR–pMHC interaction, we first compared the occupancy of salt bridges and H-bonds between SMD and force-free MD simulations in the absence of CD8. Accompanied by the rolling, the TCR leans toward the N-terminal region of the peptide. For clarity, this orientation is hereafter referred to as the distal end of the α_1_-α_2_ platform, whereas the opposite boundary flanking the peptide C-terminus is termed the proximal end. After TCR inclined to the distal end, several interfacial residue pairs on the distal end gained more chances to form interactions with TCR loops and exhibited increased occupancy under force (Fig. 9a, b). The residue D55 in β CDR2 and D31 in β CDR2 are the key residues, responsible for establishing new interactions. The force-induced rolling brings D55 and D31 close enough to form salt bridge or H-bond interactions with R75 and K146, respectively (Fig. 9c). The occupancies of H-bonds and salt bridges between D55–R75 and D31–K146 rose by ∼20% to ∼50% (Fig. 9a, b).

**Fig. 9.**
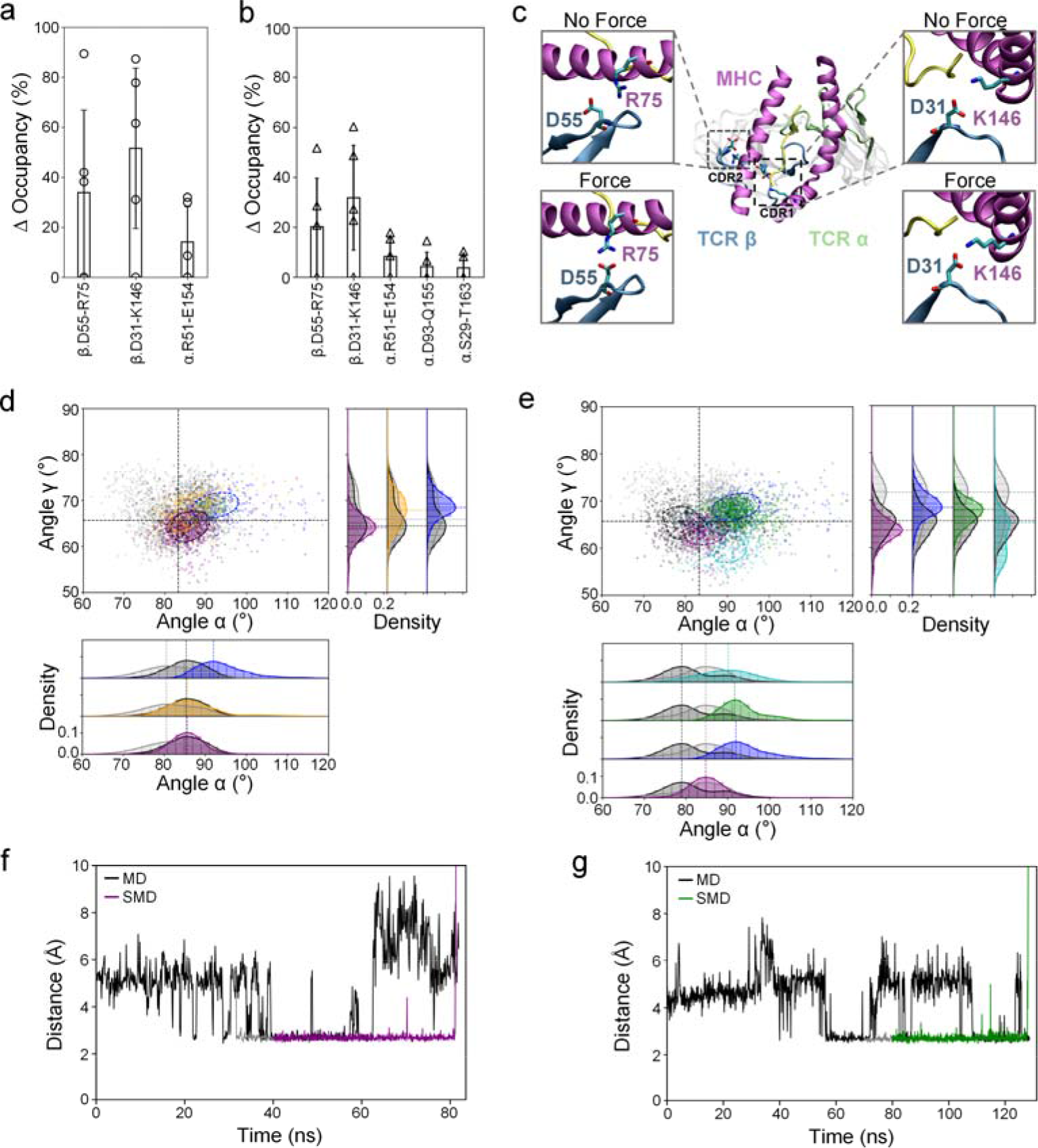
Force-induced enhancement of TCR–pMHC interfacial interactions in the absence of CD8. a, b. Occupancy increase in salt bridges (a) and hydrogen bonds (b) under force. Only residues with ΔOccupancy > 5% are presented. c. Representative snapshots of D55–R75 and D31–K146 interactions strengthened upon application of force in the CD8– system. d, e. The scatter plot in the α–γ plane (upper left) and the density histogram of rolling angle α (lower) and torsion angle γ (upper right). The dots are colored according to the interaction condition of D55–R75 (d) and D31–K146 (e). Different colors correspond to different simulation runs as indicated in Fig. 7. Black dots indicate the configurations with interaction formed in force-free MD simulation. Gray dots indicate interaction is not formed. The dashed circle highlights the density contour of the cluster representing the top 2% of the populated structures of each simulation run. Dashed lines in the density histograms indicate the peak positions of the α and γ angle distributions for the respective runs. The color code is consistent with the scatter plot. f, g. Time evolution of inter-residue distance of D55–R75 (f) and D31–K146 (g) in representative SMD trajectories. Black curves represent force□free MD trajectories, gray curves represent 10□ns force□free MD relaxation trajectories after the enlargement of the simulation box, and colored curves represent representative SMD trajectories.

To validate the correlation between the rolling and the strengthened interaction, we further produced a scatter plot of the simulated configurations in the α–γ space (Fig. 9d, e). The colored dots and black dots represent the configurations with the specific interaction formed in the MD or SMD simulations, respectively (Fig. 9d for D55–R75 and Fig. 9e for D31–K146), while the gray dots represent the configurations without the interaction formation. According to the scatter map, when the interaction of either D55–R75 or D31–K146 is formed, the TCR–pMHC exhibits higher rolling angles α. However, no clear difference is found in the torsion angles γ. Besides the two key residue pairs, several interfacial interactions between TCR α CDR residues (D93/R51/S29) and their MHC counterparts (Q155/E154/T163) were also enhanced upon application of force in the SMD simulations (Fig. S5). Consistently, scatter plots reveal that these interactions correlate with larger rolling angles α rather than torsional angles γ (Fig. S5b-d). These results suggest that force-induced rolling reorganizes the interfacial geometry, pairing the residue on both sides in a more favorable spatial arrangement.

The distance change of the key residue pairs was monitored to examine whether the interactions are stable or not (Fig. 9f, g). In the free MD simulations, occasional drops and rises in the distances were observed for both D55–R75 and D31–K146 (Fig. 9f, g), indicating that neither of the residue pairs forms stable interactions. In contrast, with force applied to the complex, the distances between these residue pairs remained below a 3.2 Å threshold before the dissociation, confirming that these salt bridges are structurally stable under load.

### 3.5 Force modulates the TCR–pMHC interaction through a different manner in the presence of CD8

Since the CD8 binding changes the force-induced dissociation pathway, making it dominated by twisting instead of rolling, the force-strengthened interaction pairs in TCR–pMHC complex may not be enhanced in the TCR–pMHC–CD8 complex. To validate this hypothesis, we analyzed the variation of TCR–pMHC interaction during the SMD simulations of the TCR–pMHC–CD8 complex. The interaction analysis reveals that the mechanical tension enhances several polar interactions (Fig. 10a, b). The most important force-enhanced interaction occurred between TCR β D55 and MHC K68, exhibiting a 30% elevation in the salt bridge occupancy upon application of force. It is worth noting that completely different sets of stabilized bonds were observed with or without CD8, suggesting CD8 engagement fundamentally alters the network of force-enhanced polar interactions (Fig. 10c and Fig. S7a). In the absence of CD8, the TCR–pMHC interface exhibits significant reorganization accompanied by the rotation of torsion and pitching during force-free MD. As a result, the distances between these key residue pairs are significantly enlarged. Even under force, these residues cannot be brought close enough for effective interactions.

**Fig. 10.**
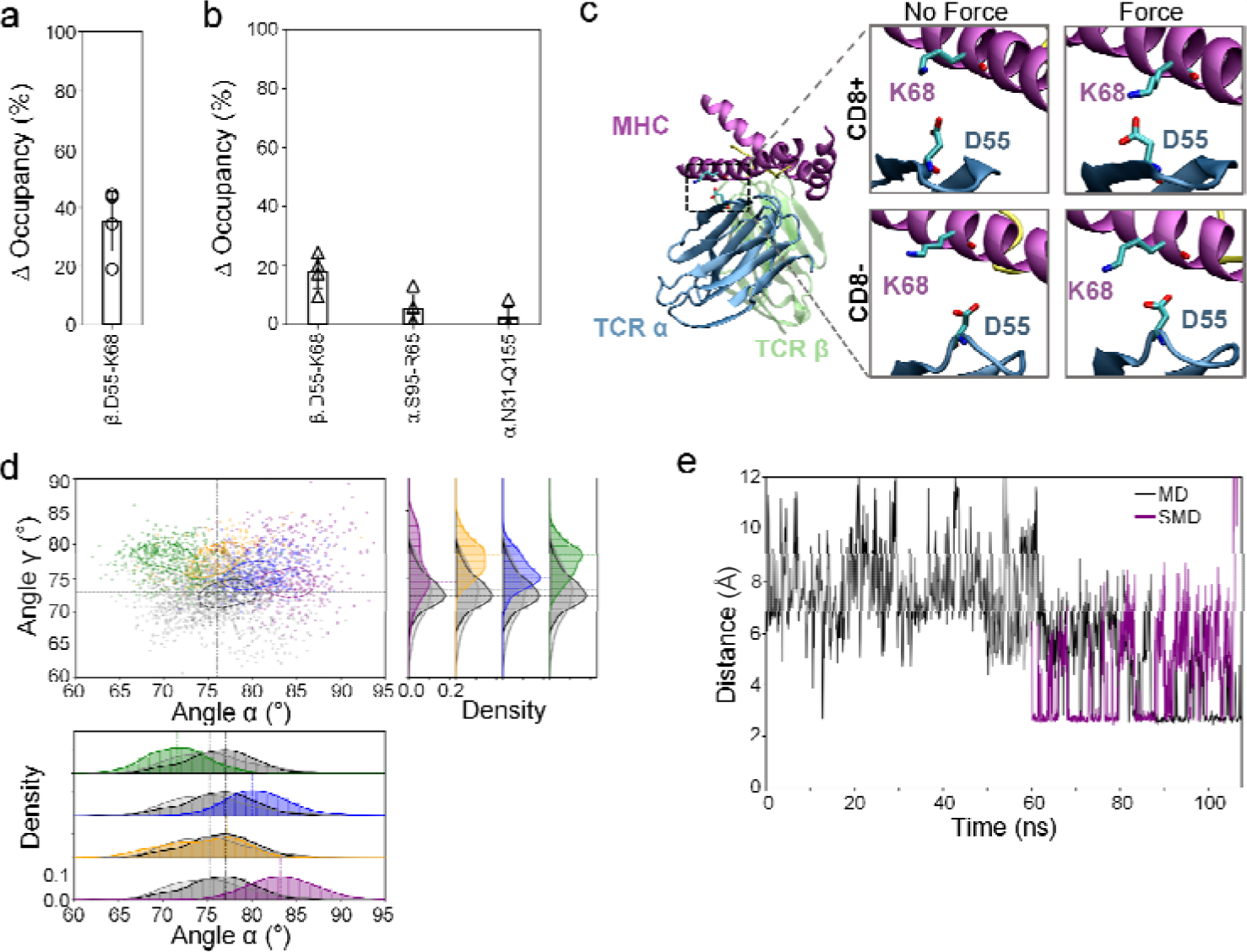
CD8-mediated enhancement of TCR–pMHC interfacial interactions induced by increased angle γ. a, b. Occupancy increase in salt bridges (a) and hydrogen bonds (b) under force. Only residues with ΔOccupancy > 5% are presented. c. Representative snapshots of D55–K68 interfacial interaction with (right) or without (left) force in the CD8– (lower) or CD8+ (upper) systems. d. The scatter plot in the α–γ plane (upper left) and the density histogram of rolling angle α (lower) and torsion angle γ (right). The dots are colored according to the interaction of D55–K68. The symbols and lines share the same definition as in Fig. 9d. e. A representative time evolution of inter-residue distance for D55–K68 in SMD simulations. Black curves represent force-free MD trajectories, gray curves represent 10-ns force-free MD relaxation trajectories after the enlargement of the simulation box, and colored curves represent representative SMD trajectories.

Furthermore, we mapped the configurations in the MD and SMD to the α–γ space and colored the dots if D55 in TCR β CDR2 forms a salt bridge with K68. Without applying force, the salt bridge formation between D55 and K68 did not affect the distribution of the rolling angle α. However, it greatly impeded the fluctuation of the torsion angle γ without changing its mean, as suggested by the superimposed but sharpened peak of the distribution of γ (Fig. 10d). With the force applied, the torsion angles of the configurations with salt bridge formed maintained a narrow distribution, with the mean of torsion shifted up by ∼5° to ∼10° (colored dots in the scatter plot and colored curve in the density histogram of Figure 10d). The force narrowed the rolling angle distribution, but the mean values did not change in any consistent pattern (Figure 10d). Force-enhanced α N31–Q155 and α S95–R65 hydrogen bonds exhibited similar shifts toward higher torsional angles γ in the CD8-containing system (Fig. S7b,c).

Without restriction from CD8, a clear twist was observed during the relaxation of the TCR–pMHC complex according to the sudden jump in the torsion angle analysis (Fig. 2e). After the rotation of twisting, the residue W96 of Vβ gets close enough to the hydrophobic pocket and has a great chance to insert in and stably anchor to the pocket (Fig. 5f). In contrast, with CD8 binding to MHC, the co-receptor stabilized the TCR–pMHC in the initial conformation. The W96 barely reaches the hydrophobic pocket. As a result, TCR–pMHC could hardly form the hydrophobic interaction between β CDR3 and the MHC α_2_ hydrophobic pocket, in the presence of CD8. However, with tension applied to the TCR–pMHC–CD8 complex, MHC twisted around TCR with the torsion angle elevated by ∼5° to ∼10°. The twisting between TCR–pMHC brought W96 to the hydrophobic pocket, allowing the hydrophobic interaction to form. Mapping the simulation conformations onto the α–γ space revealed that the force-induced insertion of W96 was closely associated with the shift toward higher torsion angles γ, whereas the rolling angle α remained broadly distributed (Fig. 11b). In the SMD simulation trajectories, the insertion of W96 to the pocket was observed under force (Fig. 11a). To quantitively investigate this interaction, the SASA and distance between β W96 and V147 were calculated. Both the distance and SASA are significantly lower under force than those in the force-free condition, and the distribution of SASA showed a single sharp peak (Fig. 11c, d). These results confirmed that in the presence of CD8, tensile force drives the twisting of TCR–pMHC which facilitates the insertion of W96 and the formation of the hydrophobic interaction. These findings indicate that CD8 not only shifts the dissociation trajectory but also reshapes the TCR–pMHC interaction landscape under force.

**Fig. 11.**
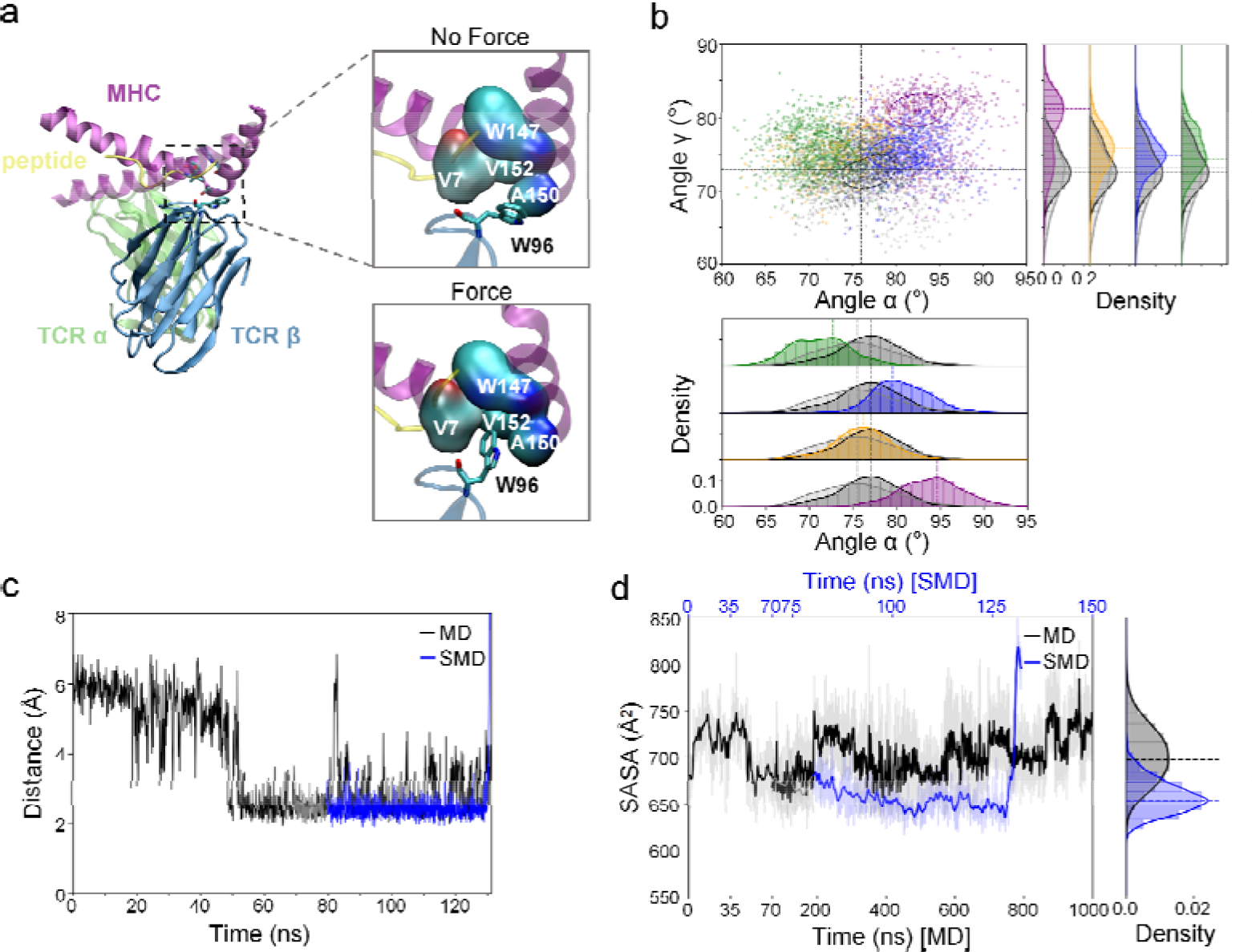
Occupancy of the hydrophobic pocket driven by angle and its stability under force. a. Representative snapshots of the hydrophobic pocket (formed by the peptide and the MHC α_2_ helix) occupied by TCR residue W96 upon application of force in the CD8+ system. b. The scatter plot in the α–γ plane (upper left), the density histogram of rolling angle α (lower), and torsion angle γ (right). The dots are colored according to whether residue W96 enters the hydrophobic pocket. The symbols and lines share the same definition as in Fig. 9d. c. A representative time-course of the distance between pocket center (W147) and W96 in the SMD trajectory. d. Time-course (left) and density histogram (right) of the SASA values for the hydrophobic pocket in the representative SMD trajectory. Black curves represent force-free MD trajectories, gray curves represent 10□ns force-free MD relaxation trajectories after the enlargement of the simulation box, and colored curves represent representative SMD trajectories.

## 4. Discussion

Previous force-free MD studies have suggested that fluctuations of the antigenic peptide and the MHC binding groove can alter key interfacial contacts and influence the stability of productive TCR–pMHC binding configurations (29,57). Wan et al. constructed a TCR–pMHC–CD4 complex embedded in cell membrane and simulated the system to investigate the free dynamics characteristics (58). Lately, Rollins et al. performed SMD simulation on the CD3–TCR–pMHC–CD4 complex and studied the interaction energy and H-bond changes during the dissociation of pMHC from the complex (59). Herein, we performed free MD or SMD simulations on the TCR–pMHC or TCR–pMHC–CD8 complex embedded in cell membrane. Under force-free conditions, the TCR–pMHC complex without CD8 is less stable, with significant fluctuations in intermolecular rotations accompanied by the redistribution of key contact residues at the recognition interface. In contrast, in the trimolecular complex, CD8 stabilizes the MHC α_3_ domain and restricts the intermolecular rotation of the complex. Both the rolling and twisting angles are limited in a narrow range with a single-peak distribution (Fig. 2c-f). As a result, the interacting residues are more stable than those in the bimolecular complex (Fig. 5a,b). The results suggest that CD8 binding to the MHC α_3_ domain provides an additional structural constraint, narrows the accessible conformational space of the TCR–pMHC complex, stabilizes key recognition contacts at the interface under force-free conditions, and thereby converts the highly dynamic TCR–pMHC interface into a more restricted and stable recognition state. Earlier experimental work showed that CD8 promotes TCR–pMHC recognition by forming cooperative TCR–pMHC–CD8 trimolecular interactions and by reducing dissociation rates or prolonging bond lifetimes (24,25). The CD8 binding-induced stabilization of the TCR–pMHC provides a structural explanation for these observations.

The mechanical environment of the immunological synapse plays a vital role in T cell activation. A hallmark of this process is the catch-bond signature of the TCR–pMHC complex, which allows agonist-bound receptors to resist force-induced dissociation and maintain a stable signaling platform (23). In our SMD simulations, the two systems exhibit distinct response to the tensile force. The TCR rolls on the pMHC platform under mechanical loading in the TCR–pMHC complex. The observed tension-driven rolling aligns with previous reports identifying pMHC rotation as a key mechanical event that underlies catch-bond behavior during agonist recognition (22). Notably, in the TCR–pMHC–CD8 complex the rolling is impaired but twisting of TCR–pMHC is remarkable under tension. This indicates a dissociation pathway shifts from the rolling dominated pattern to the twisting dominated pattern with CD8 binding. Moreover, the sliding of CD8 from MHC α_3_ domain to MHC α_1_-α_2_and occasionally inserting into the TCR–MHC interface establishes a new trimolecular interaction network. This suggests that CD8 regulates the structural organization and remodeling of the interface during force loading. Furthermore, coordinating with CD8, tensile force-induced twisting brings the hydrophobic surface on TCR and pMHC close enough to form an additional interaction. Taken together, the TCR–pMHC–CD8 complex can maintain a more stable binding state under mechanical stress in the immunological synapse. Given that the prolonged binding lifetime provides sufficient time for downstream signaling events suggested by the kinetic proofreading model (60), CD8 and tensile force cooperatively facilitate the triggering the T cell activation.

While this study provides atomic-level insights into how CD8 modulates the conformational dynamics and dissociation behavior of the TCR–pMHC complex, several limitations should be acknowledged. First, the CD3 signaling subunits are not included in the present simulations. Previous studies have shown that antigen recognition-induced conformational and dynamical changes at the TCR–pMHC interface can affect the TCR constant and transmembrane regions, thereby modulating the TCR–CD3 signaling complex (32,61). Therefore, whether the CD8-mediated structural rearrangements identified here can be further coupled to CD3 activation and downstream signaling remains unclear. Second, this study focuses on a single TCR–pMHC complex, whereas natural TCR repertoires exhibit enormous diversity. Future studies involving additional TCR–pMHC systems will be required to further evaluate how the molecular mechanisms identified here may vary across different antigen recognition contexts.

## 5. Conclusion

In this study, we performed all-atom MD and SMD simulations on TCR–pMHC or TCR–pMHC–CD8 complexes in a membrane environment to examine how CD8 and force regulate the TCR–pMHC interaction. In the force-free condition, the binding of CD8 to the MHC α_3_ domain introduces strong constraints not only to MHC molecules but also to the TCR–pMHC interface. Pulling force applied to the TCR–pMHC complex induces rolling of pMHC relative to TCR, which leads to additional interaction pairs between the interface. However, the binding of CD8 alters the dissociation pathway, in which a consistent twisting is found in all the simulation runs. In summary, this study revealed the mechanisms of CD8/force regulation of TCR–pMHC interaction at the atomic scale, deepening our understanding of the T cell antigen recognition.

## Acknowledgments

This work is supported by grants from the National Natural Science Foundation of China (12272216 and 12172204).

