## Supplementary Figures for "CD8-mediated organization of the TCR–pMHC interface shapes its force response and dissociation pathways"

Jing Li^1^, Zhenhai Li^1^*

Zhenhai Li

.

**This PDF file includes:**

Figures S1 to S7

Legends for Movies S1 to S7

**Other supporting materials for this manuscript include the following:**

Movies S1 to S7

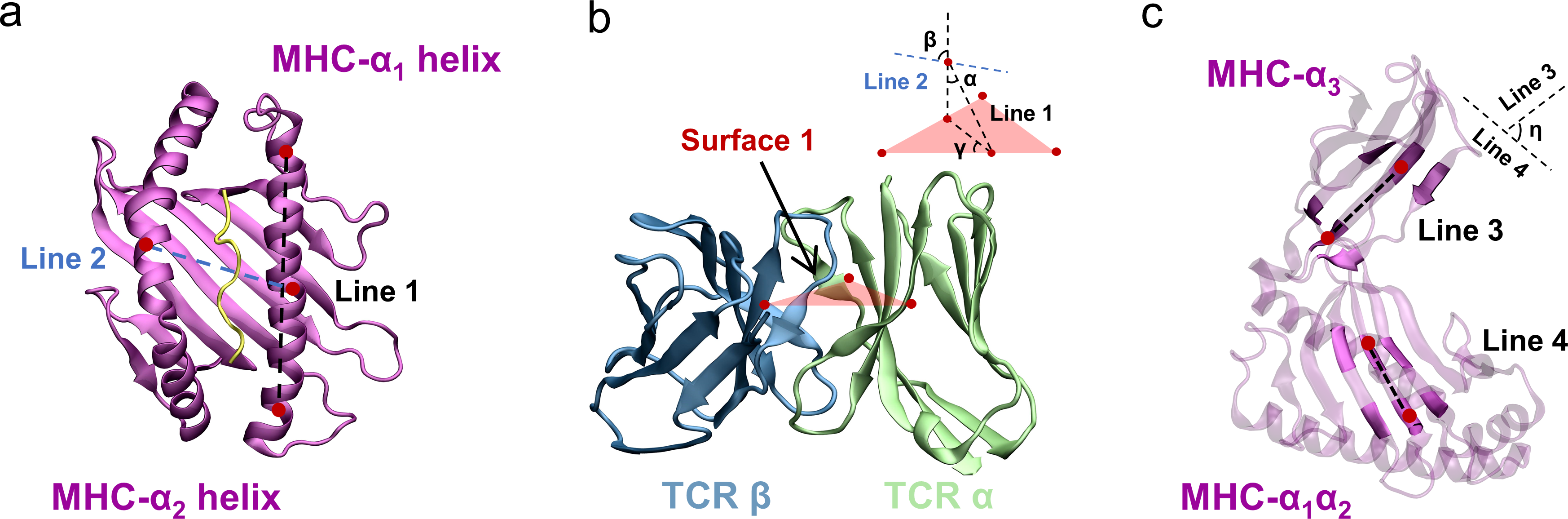

Figure S1. Geometric definitions of the relative orientation of TCR–pMHC and the internal conformation of MHC. a, b. Definitions of the intermolecular orientation of the MHC relative to the TCR. Line 1 and Line 2 represent reference vectors of the MHC, and Surface 1 is the reference plane constructed from the TCR variable domains. Based on these geometric elements, the rolling angle α, pitching angle β, and torsion angle γ are defined. c. Geometric definitions of MHC hinge angle η. Line 3 and Line 4 represent characteristic vectors of the α_1_-α_2_ platform and the α_3_ domain, respectively, and the hinge angle η between them quantifies the relative conformational change of the two domains.

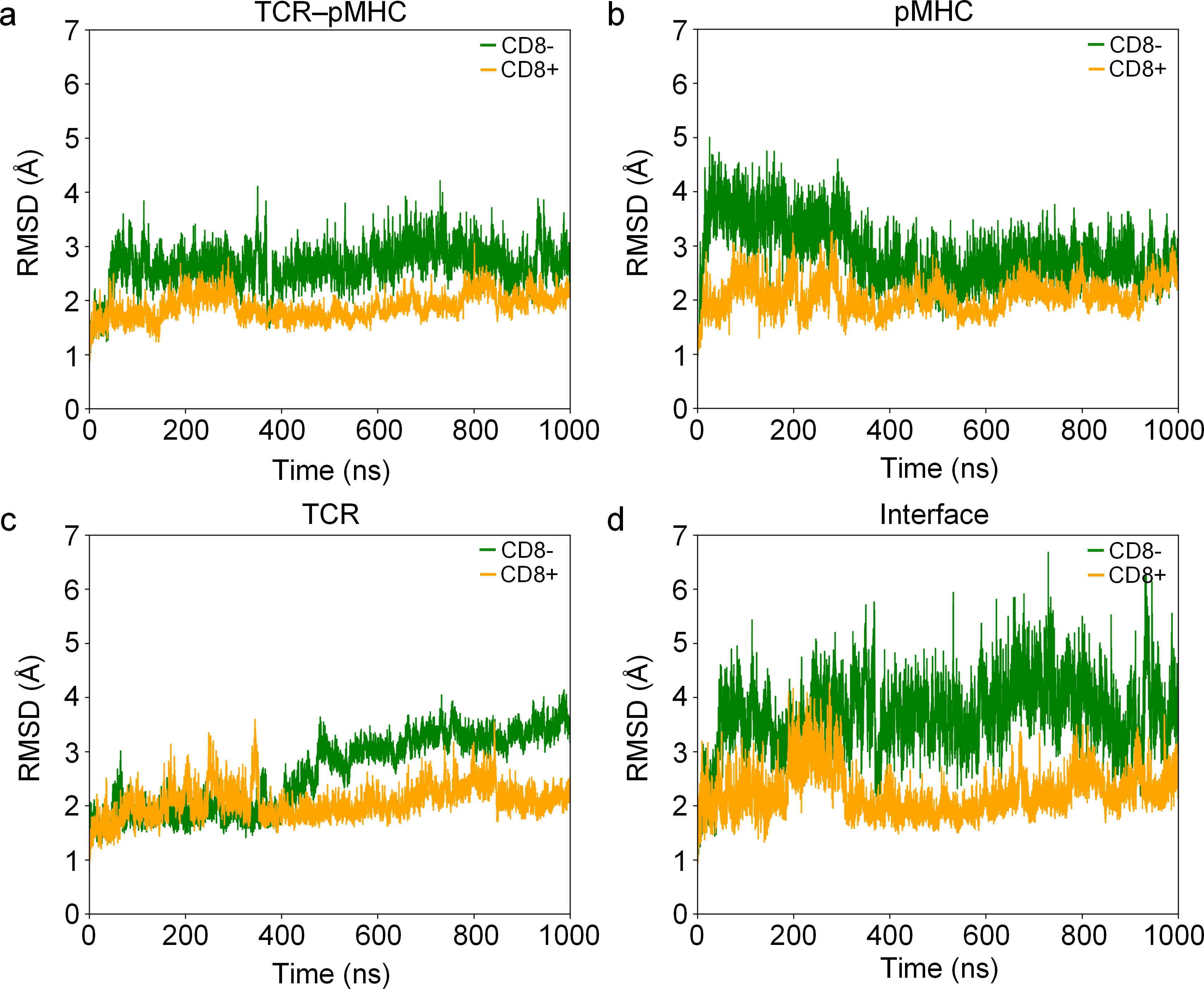

Figure S2. Structural stability analysis of the TCR–pMHC complexes. Root Mean Square Deviation (RMSD) profiles of backbone atoms for the TCR–pMHC (green) and TCR–pMHC–CD8 (orange) systems during 1 μs force-free molecular dynamics simulations. RMSD was calculated for the entire extracellular TCR–pMHC complex (a), pMHC (b), the extracellular domain of TCR (c), and the TCR–pMHC binding interface (d).

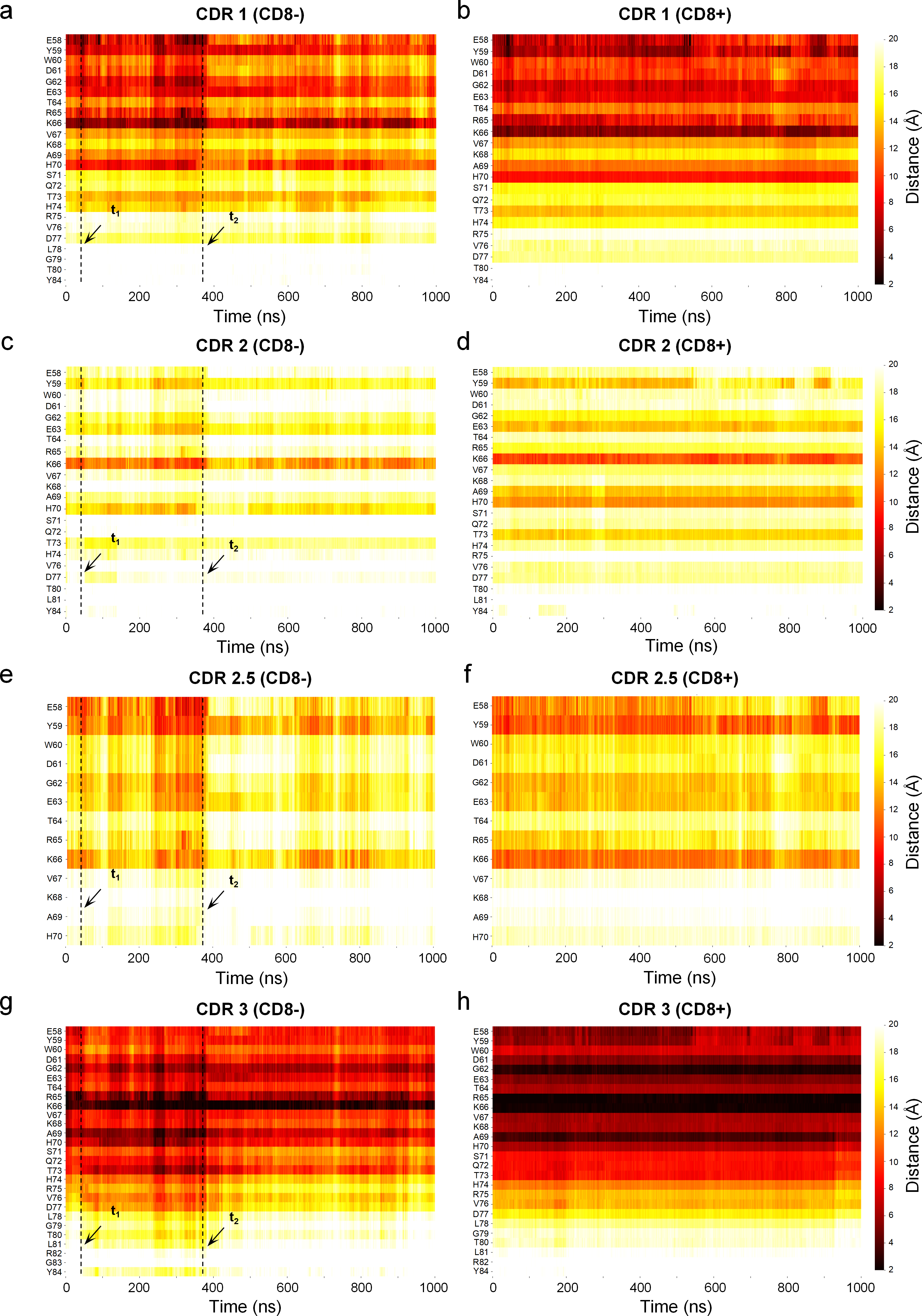

Figure S3. Time course of the contacts between TCR α CDR loops and the MHC α1 helix during Force-free MD simulations. a-h. Time courses of minimum distances between MHC α1 helix residues and TCR α CDR1 (a, b), CDR2 (c, d), CDR2.5 (e, f), and CDR3 (g, h) loops in the TCR–pMHC system (a, c, e, g) and TCR–pMHC–CD8 system (b, d, f, h). Time points t1 and t2 mark interface sliding events associated with γ angle shifts.

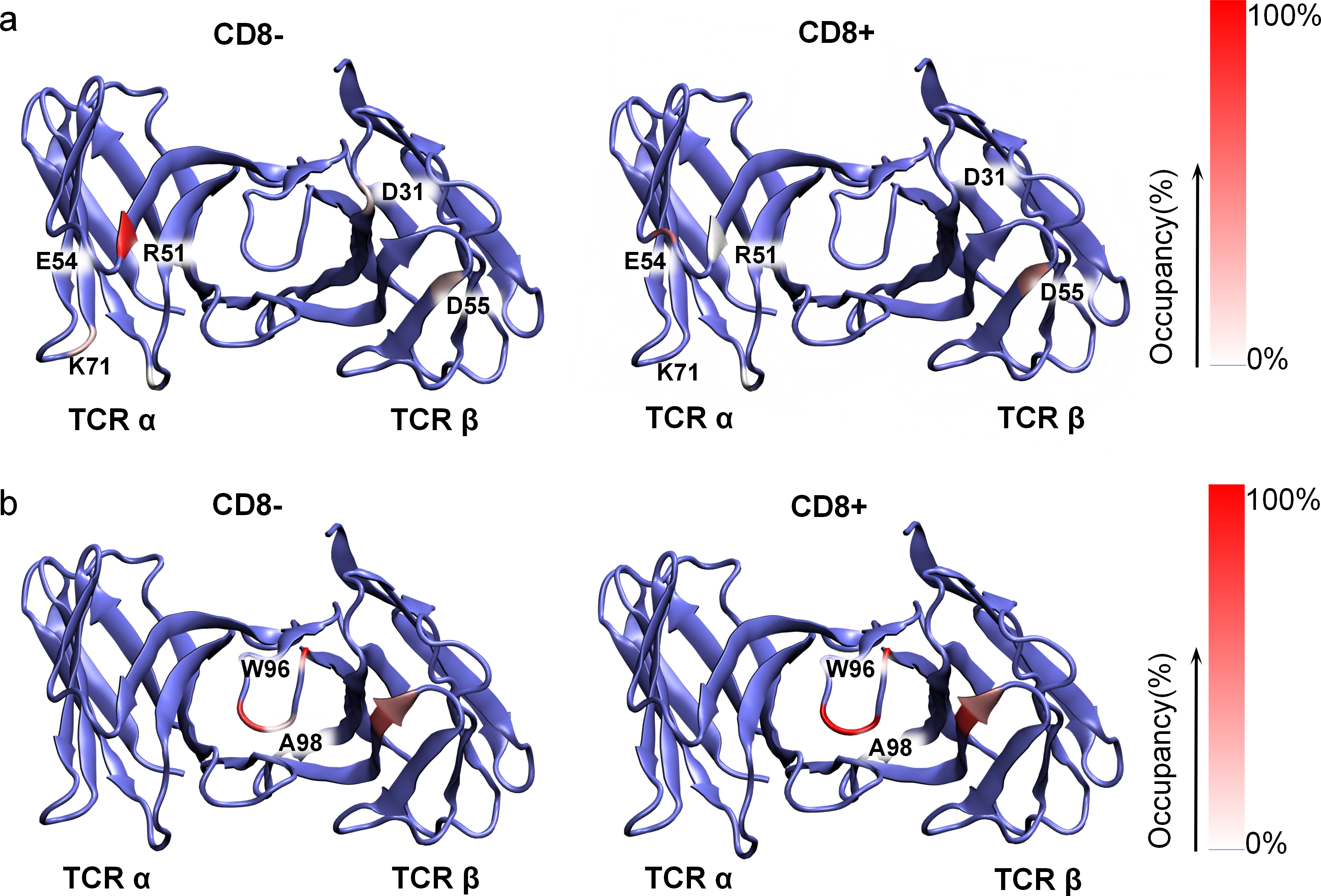

Figure S4. CD8-mediated rearrangement of recognition hotspots. a, b. The heatmaps of salt bridge (a) and hydrophobic (b) interaction occupancy on the TCR V domains in the absence (left) or presence of CD8 (right).

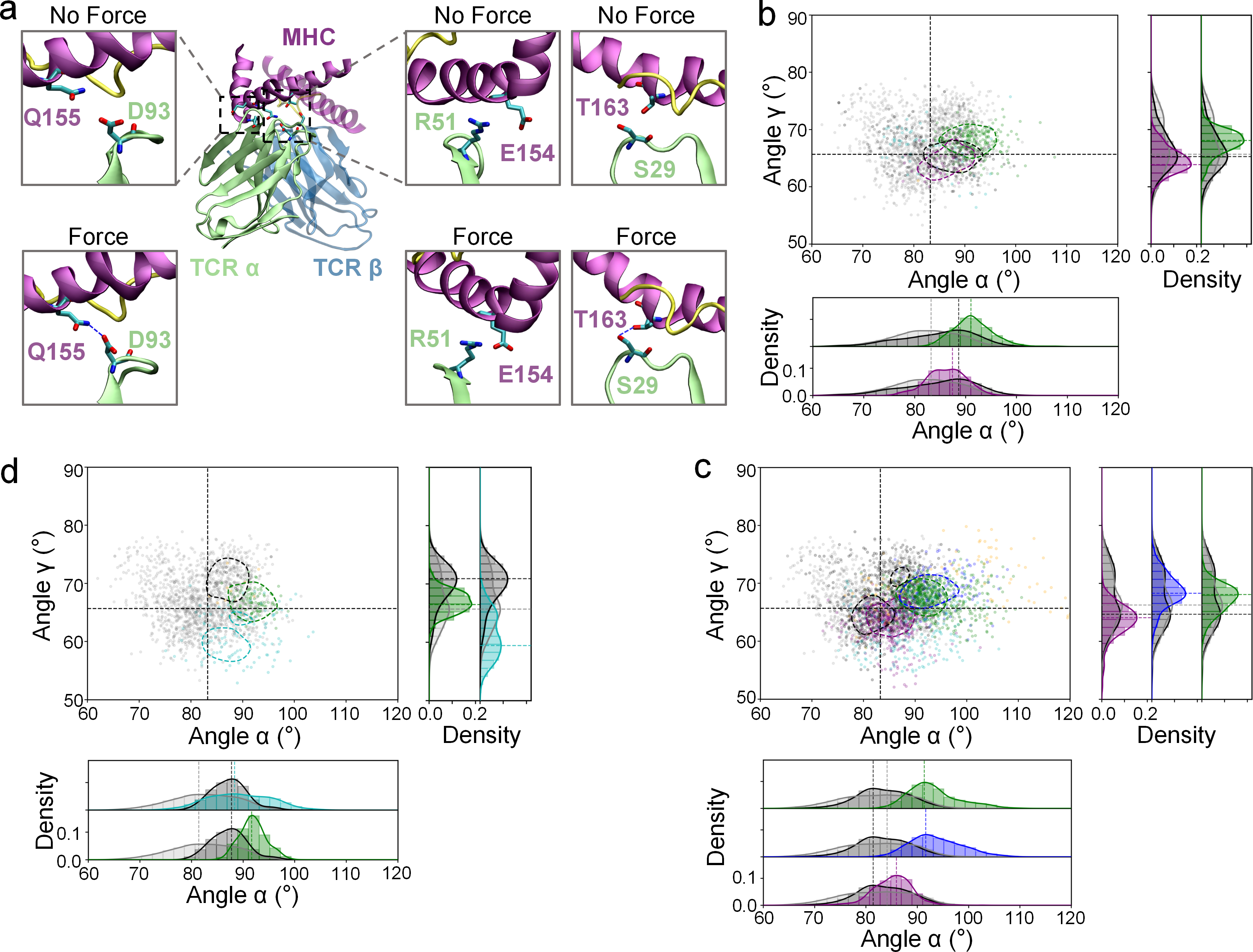

Figure S6. Additional force-enhanced interfacial interactions in the TCR–pMHC system. a. Representative snapshots of D93–Q155, R51–E154, and S29–T163 interfacial interactions strengthened upon application of force in the CD8– system. Hydrogen bonds are shown as blue dashed lines. b-d. The scatter plot in the α–γ plane (upper left) and the density histogram of rolling angle α (lower) and torsion angle γ (upper right). The dots are colored according to the interaction condition of D93–Q155 (b), R51–E154 (c), and S29–T163 (d). The symbols and lines share the same definition as in Fig. 9d.

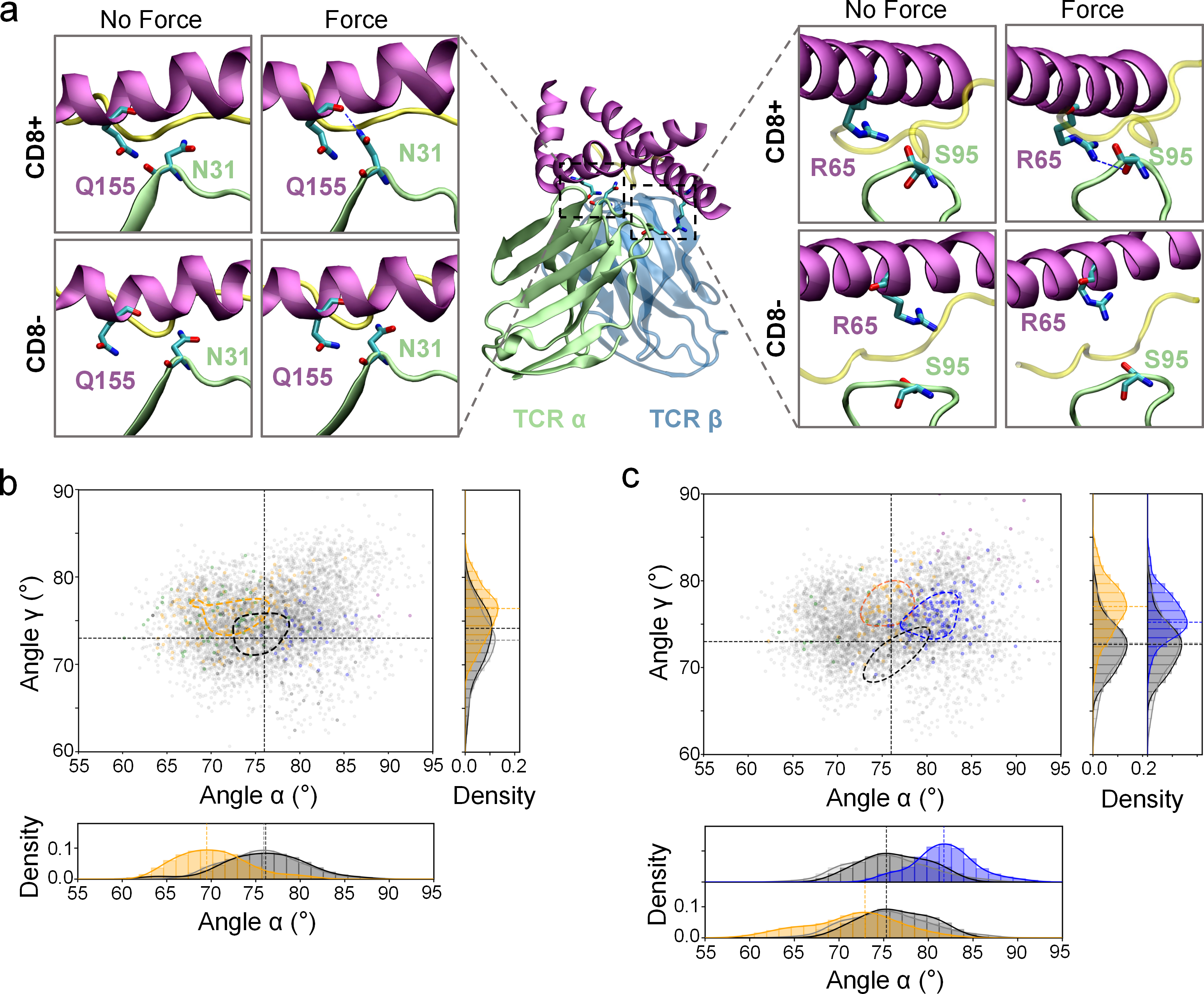

Figure S7. Additional examples of CD8-mediated enhancement of TCR–pMHC interfacial interactions. a. Representative snapshots of N31–Q155 and S95–R65 interactions under force-free MD and SMD simulation, comparing TCR–pMHC (lower) and TCR–pMHC–CD8 (upper) systems. Hydrogen bonds are shown as blue dashed lines. b, c. The scatter plot in the α–γ plane (upper left) and the density histogram of rolling angle α (lower) and torsion angle γ (upper right). The dots are colored according to the interaction condition of N31–Q155 (b) and S95–R65 (c). The symbols and lines share the same definition as in Fig. 9d.

**Video S1.** 1000-ns force-free MD simulations of TCR–pMHC (left) and TCR–pMHC–CD8 (right) systems.

**Video S2.** Changes in rolling angle α during 1000-ns force-free MD simulations of TCR–pMHC (left) and TCR–pMHC–CD8 (right) systems. Trajectories were aligned to the TCR variable region for visualization. Only the TCR variable region, MHC α_1_-α_2_ domain, and peptide are shown.

**Video S3.** Changes in pitching angle β during 1000-ns force-free MD simulations of TCR–pMHC (left) and TCR–pMHC–CD8 (right) systems. Trajectories were aligned to the TCR variable region for visualization. Only the TCR variable region, MHC α_1_-α_2_ domain, and peptide are shown.

**Video S4.** Changes in torsion angle γ during 1000-ns force-free MD simulations of TCR–pMHC (left) and TCR–pMHC–CD8 (right) systems. Trajectories were aligned to the TCR variable region for visualization. Only the TCR CDR loops, MHC α_1_-α_2_ domain, and peptide are shown.

**Video S5.** Changes in hinge angle η during 1000-ns force-free MD simulations of TCR–pMHC (left) and TCR–pMHC–CD8 (right) systems. Trajectories were aligned to the MHC α_1_-α_2_ domain for visualization. Only the whole MHC molecule is shown.

**Video S6.** Five independent SMD trajectories of the TCR–pMHC system.

**Video S7.** Four independent SMD trajectories of the TCR–pMHC–CD8 system.
